# Prefrontal default-mode network interactions with posterior hippocampus during exploration

**DOI:** 10.1101/2025.03.12.642890

**Authors:** Andrew E. Papale, Vanessa M. Brown, Angela M. Ianni, Michael N. Hallquist, Beatriz Luna, Alexandre Y. Dombrovski

## Abstract

Hippocampal maps and ventral prefrontal cortex (vPFC) value and goal representations support foraging in continuous spaces. How might hippocampal-vPFC interactions control the balance between behavioral exploration and exploitation? Using fMRI and reinforcement learning modeling, we investigated vPFC and hippocampal responses as humans explored and exploited a continuous one-dimensional space, with out-of-session and out-of-sample replication. The spatial distribution of rewards, or value landscape, modulated activity in the hippocampus and default network vPFC subregions, but not in ventrolateral prefrontal control subregions or medial orbitofrontal limbic subregions. While prefrontal default network and hippocampus displayed higher activity in less complex, easy-to-exploit value landscapes, vPFC-hippocampal connectivity increased in uncertain landscapes requiring exploration. Further, synchronization between prefrontal default network and posterior hippocampus scaled with behavioral exploration. Considered alongside electrophysiological studies, our findings suggest that locations to be explored are identified through coordinated activity binding prefrontal default network value representations to posterior hippocampal maps.

**Significance Statement:** The ventral prefrontal cortex represents goals and values, while the hippocampus contains maps of physical and abstract spaces. In recent years, neuroscientists have sought to understand how hippocampal-prefrontal interactions help us to solve complex problems such as deciding whether to exploit known rewards or explore in search of better alternatives. Using functional magnetic resonance imaging and reinforcement learning modeling, we examine these interactions as humans explore and exploit an environment with a complex distribution of rewards or value landscape. We describe how this value landscape modulates activity and connectivity between prefrontal cortex and hippocampus, controlling the balance between exploration and exploitation.

## Introduction

In uncertain environments, animals face the dilemma of whether to *exploit* known good options or *explore* potentially better alternatives. Behavior shifts from exploration to exploitation as valuable options are discovered. In environments with a few discrete options mammalian striatum and amygdala can resolve this dilemma efficiently (Costa et al., 2019), but more complex spaces require hippocampal cognitive maps (Johnson et al., 2012; Mehlhorn et al., 2015; Dombrovski et al., 2020) and flexible reward representations in ventral prefrontal cortex (vPFC; Domenech et al., 2020; Trudel et al., 2021).

Locations of desirable outcomes can be predicted by a cognitive map integrating available cues with spatial memory. The hippocampus creates value-laden maps based on trajectories through space (Stachenfeld et al., 2017; Dombrovski et al., 2020). These maps contain more fine-grained representations in posterior/dorsal divisions and reward- and context-based representations in anterior/ventral divisions (Royer et al., 2010; Biane et al., 2023). Moreover, the hippocampus replays trajectories (Skaggs and McNaughton, 1996; Liu et al., 2019; Schuck and Niv, 2019) or considers future goals online during deliberation (Johnson and Redish, 2007; Papale et al., 2012; Redish, 2016). vPFC, in turn, estimates option values based on current needs and goals (Padoa-Schioppa, 2007; Bartra et al., 2013; Tang et al., 2021). We have found that human anterior hippocampus represents spatially structured values, and these representations scale with successful exploitation (Dombrovski et al., 2020). However, how interactions between various hippocampal and vPFC subregions may support exploration and exploitation remains unclear.

Hippocampal-vPFC interactions are facilitated by long-range synchronization of local field potentials (Buzsáki, G., 2006). Consistent with behavioral evidence, offline delta synchronization (Schultheiss et al., 2020) – linked to sharp wave ripples in hippocampus – promotes memory consolidation and retrieval of contextually relevant information (Buzsáki, 2015; Joo and Frank, 2018; Tang and Jadhav, 2019), potentially including option values (Weilbächer and Gluth, 2016; Schmidt and Redish, 2021). Additionally, theta synchronization during directed attention conveys hippocampal spatial representations to the vPFC (Jones and Wilson, 2005; Gluth et al., 2015; Weilbächer and Gluth, 2016).

Value-related information differentially modulates vPFC subregions that fall into three large-scale connectivity networks: the ventral limbic network (LIM), the default mode network (DMN) and the frontoparietal control network (CTR). The value of the chosen or best available option positively modulates medial and central orbitofrontal and subcallosal LIM as well as peri-callosal and fronto-polar DMN (Hare et al., 2008; Bartra et al., 2013; Smith et al., 2014; Lopez-Gamundi et al., 2021; Knudsen and Wallis, 2022). By contrast, lateral OFC regions such as 11/47 belonging to CTR are negatively modulated by expected value (Ursu et al., 2008), suggesting competition among similarly-valued options for action selection. Connectivity between the putative rodent analogue of LIM (infralimbic PFC and ventral orbitofrontal cortex, OFC) and ventral hippocampus appears necessary for exploitation of learned values (Chudasama et al., 2012). In addition, interactions between hippocampus and more dorsal rodent prelimbic PFC (putatively homologous to human DMN; Lu et al., 2012) may govern exploitation (Laureiro-Martínez et al., 2015) and exploration (Morici et al., 2022). Given the diverging value responses of the CTR subregion, we reasoned that it would serve as an informative comparison when examining activity and hippocampal connectivity of DMN and LIM vPFC.

We therefore investigated (1) how spatially structured value functions or value landscapes modulate activity of the human vPFC (DMN, LIM, and CTR), as well as vPFC connectivity with anterior vs. posterior hippocampus, and (2) how such connectivity relates to behavioral exploration. To address these aims, we examined fMRI BOLD activity during exploration and exploitation of a continuous one-dimensional space. We inferred the value distribution over the space using our previously validated information-compressing RL model of adaptive transition from exploration to exploitation (Hallquist and Dombrovski, 2019). Multi-level analyses of event-locked deconvolved BOLD signal (Dombrovski et al., 2020) enabled us to characterize value encoding across within-trial timepoints, during online navigation, outcome receipt and offline processing seconds after an outcome. We replicated key findings out-of-session and out-of-sample.

## Materials and Methods

### PARTICIPANTS – EXPERIMENT 1

Participants were 71 individuals aged 14-30 (mean +/- SD: 21±5 years; 37 female, 34 male) with no history of neurological disorder, brain injury, or developmental or psychiatric disorder (self or first-degree relatives). Participants or their legal guardians provided informed consent prior to participation. Procedures complied with the Code of Ethics of the World Medical Association (1964 Declaration of Helsinki) and were approved by the Institutional Review Board of the University of Pittsburgh (protocol PRO10090478). Participants were compensated $75 for this experiment.

### PARTICIPANTS – EXPERIMENT 2

Participants were psychiatrically and neurologically healthy individuals (N=43) age 49-80 (mean +/- SD: 63±8 years; 23 female, 20 male. These data were drawn from the healthy comparison group of a study of suicidal behavior in the second half of life.

Participants were compensated $150 for completing the experiment and other studies. Experimental procedures for this study complied with Code of Ethics of the World Medical Association (1964 Declaration of Helsinki) and the Institutional Review Board at the University of Pittsburgh (STUDY19030288).

### REGIONS OF INTEREST: PREFRONTAL CORTEX AND HIPPOCAMPUS

We analyzed 17 subdivisions of vPFC, encompassing medial and lateral OFC, frontal pole, and vmPFC/pericallosal ACC. These were selected from the 7-network 200-region parcellation of (Schaefer et al., 2018; Fig. 1A). Following this parcellation, we assigned regions to one of three networks: the Default Mode (DMN, orange), the Frontoparietal (Cognitive) Control (CTR, yellow), or the Limbic (LIM, blue). Sixteen of these regions were divided into symmetric left-right pairs and labeled using the Eickhoff-Zilles Cytoarchitectonic MPM atlas 2.2 in AFNI (Fig. 1C; (Eickhoff et al., 2005, 2007; Henssen et al., 2016). To preserve the left-right symmetry, the 17th region, right frontopolar BA 10 (fp10 region 161, MNI x=13, y=64, z=-14) was moved from the limbic network to the DMN. Our sensitivity analyses showed that this change had no qualitative impact on the results in Experiment 1 for Fig. 4B, 5B, and Fig. 6C,E,G (Fig. S4, Fig. S5). The human frontal DMN expands dorsally, thus we included region dorsal BA 10 (d10 region 89 left MNI x=-7, y=52, z=11, d10 region 194 right MNI x=7, y=52, z=10) in our analyses.

**Fig. 1:**
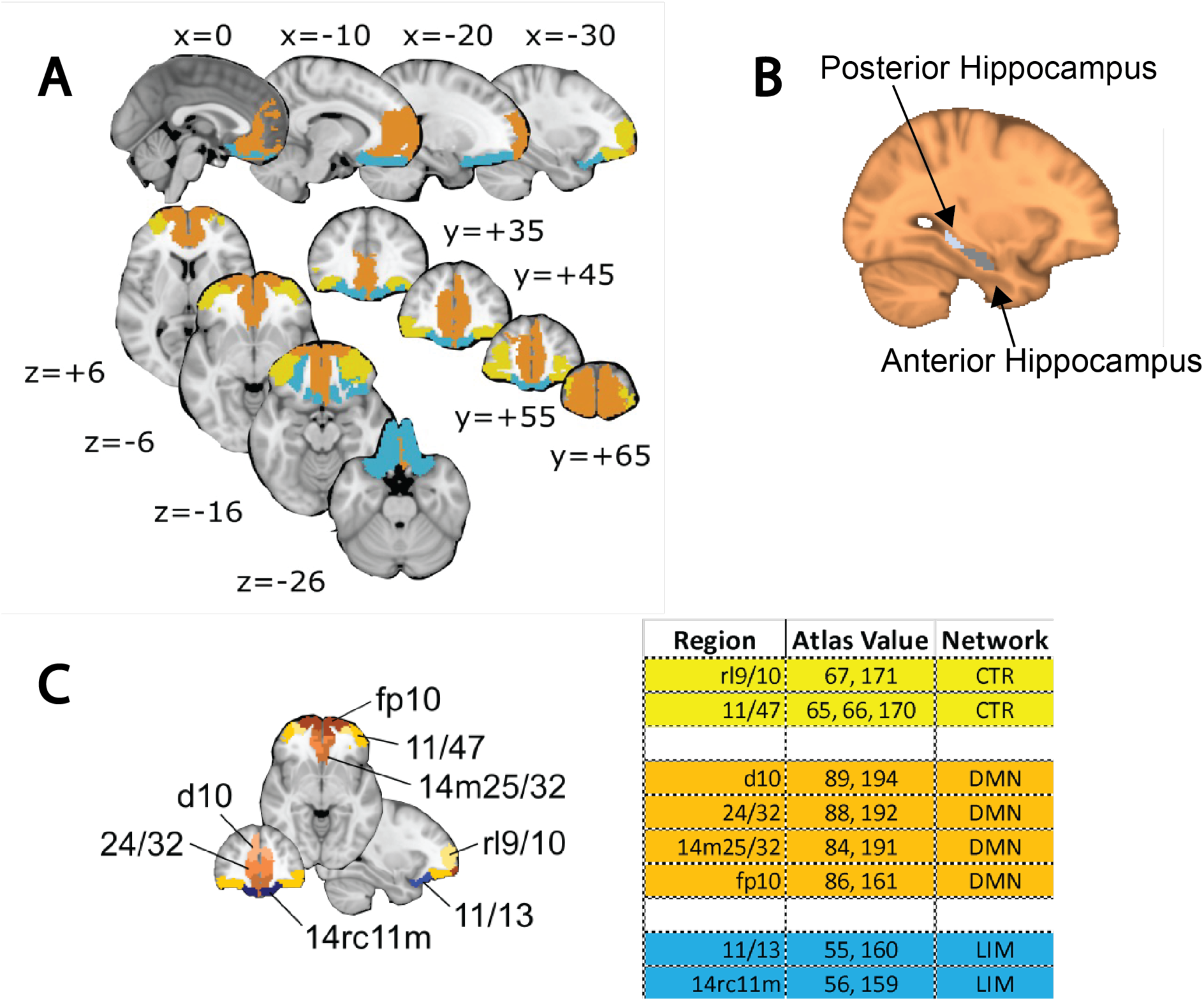
vPFC mask and hippocampus mask. **A.** The vPFC mask consists of 17 regions grouped into 3 resting-state functional networks as in Yeo et al., 2011, the Default Mode Network (DMN; orange), the Control Network (CTR; yellow), and the Limbic Network (LIM; blue). **B.** The bilateral hippocampal mask is from the Harvard-Oxford subcortical atlas. The posterior 1/3 of the hippocampus was grouped into the Posterior Hippocampus region (PH; light gray) and the anterior 2/3 of the hippocampus was grouped into the Anterior Hippocampus region (AH; dark gray). **C.** The vPFC mask is comprised of 17 parcels of the Schaefer 2018 mask derived from resting state functional connectivity. We labeled these parcels in AFNI (Cox, 1996) based on approximate overlap with the Eickhoff-Zilles Cytoarchitectonic MPM atlas 2.2. Because adjacent regions of the DMN in vPFC included what is clearly dorsal PFC, we included parcels d10 (peach shading), although it is not strictly vPFC. The vPFC mask was grouped by rough left-right symmetry into 8 distinct groups. Models were run separately with each region as a dependent variable to complement the main analyses separating vPFC into three resting state networks (Frontoparietal Control Network - CTR; yellow, Default Mode Network - DMN; orange, and Limbic Network - LIM, blue) in the 7-network solution. Since the parcellation is not strictly symmetric, one of the groups is not a pair (11/47).

The bilateral hippocampal mask was derived from the Harvard-Oxford subcortical atlas in MNI152 space (https://identifiers.org/neurovault.collection:262) and resampled to 2.3mm (Experiment 1) voxels or 3.1mm (Experiment 2) voxels to match the native resolution of the BOLD data. To delineate anterior (AH) vs. posterior hippocampus (PH), the postero-anterior long axis was defined as described previously (Dombrovski et al., 2020) with the posterior 1/3 comprising PH and anterior 2/3 comprising AH (Fig. 1B).

### BEHAVIORAL TASK – EXPERIMENT 1

On the clock task (Moustafa et al., 2008), participants were shown a clock face and asked to make a single choice per trial, stopping a dot which revolves clockwise around a central stimulus in 4s. The choice led to immediate feedback indicating points earned on that trial, displayed for 0.9s (Fig. 2A). Unknown to participants, earnings were controlled by one of four difficult reinforcement contingencies with probability-magnitude tradeoffs, in 50-trial runs. Thus, one learned that certain locations on the clock yielded higher average rewards. Two reinforcement contingencies were learnable: *increasing expected value* (IEV), where later response times yielded higher average payoffs, vs. *decreasing expected value* (DEV). In IEV and DEV, as well as one unlearnable condition, *constant expected value* (CEV), reward probabilities decreased while magnitudes decreased throughout the interval. In a final, unlearnable *constant expected value–reversed* (CEVR) condition, the probability-magnitude tradeoff was reversed. If the participant failed to make a response within 4s (360 degrees) they received zero points (Fig. 2B). Subjects consistently chose longer response times in the IEV condition (green), and shorter response times in the DEV condition (orange) in Experiment 1 (Fig. 2C) both in the fMRI (left panel) and out-of-session MEG replication (center panel) sessions.

**Fig. 2.**
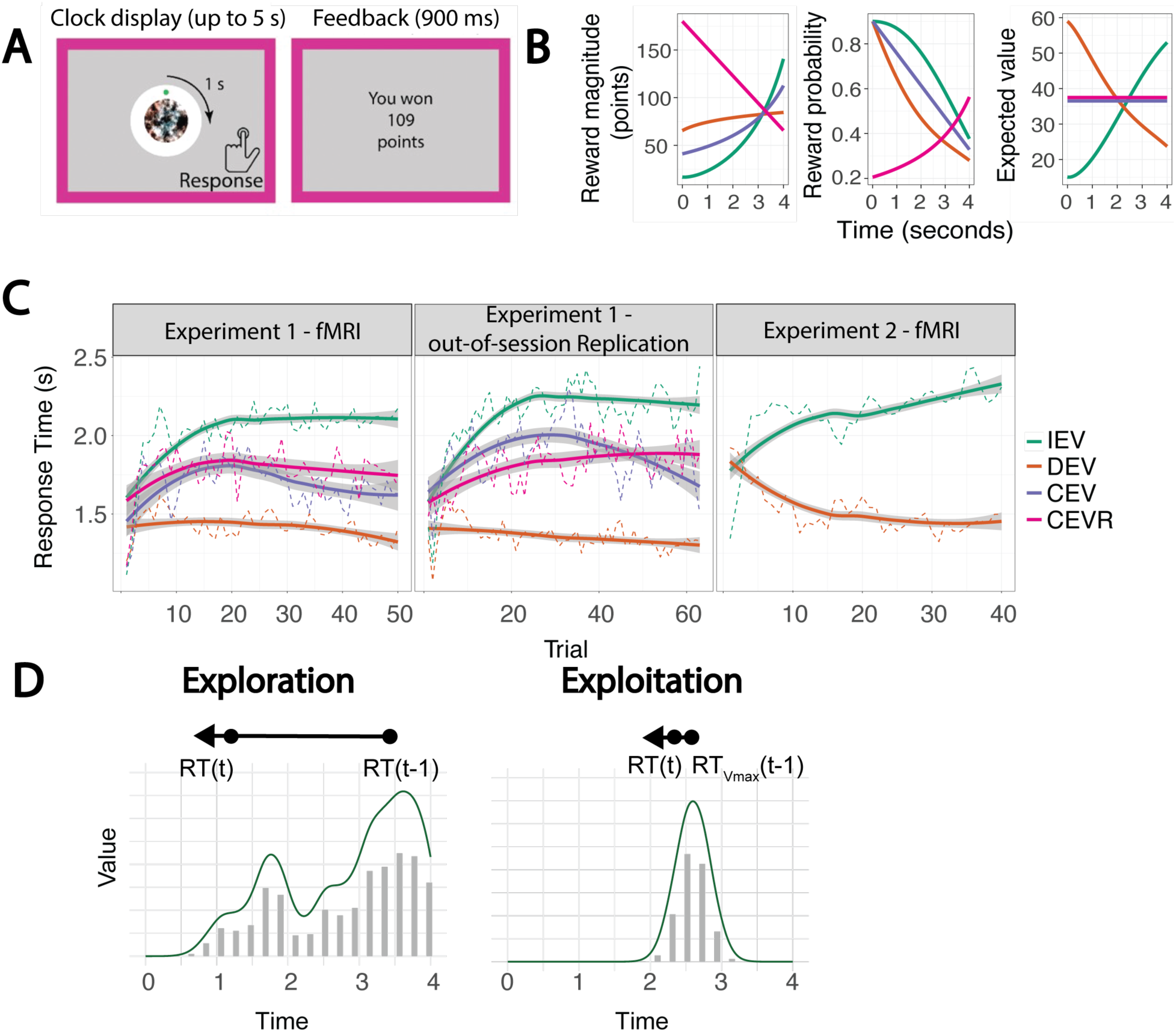
Clock task and operationalizing exploration vs. exploitation. **A.** Each trial on the clock task begins with a green dot that rotates clockwise from the 12 o’clock position around a pseudo-clock face with a central image (here a phase-scrambled image) in 4s (Experiment 1) or 5s (Experiment 2). On each trial, participants press a button to stop the green dot to earn points in each time-varying contingency. Pressing the button immediately triggers 900ms of feedback where the reward associated with that choice is displayed. **B.** Over the course of a session, the participant learns about the underlying reward contingencies and adjusts their responses accordingly. There are 4 contingencies on the clock task: increasing expected value (IEV, green), decreasing expected value (DEV, orange), constant expected value (CEV, blue) and constant expected value reversed (CEVR, pink). During IEV trials (green) subjects learn that responding later in the interval results in a higher overall payoff (greater magnitude but lower probability). In CEV (blue) and CEVR (magenta) trials, the expected value (reward probability times reward magnitude) is constant across the interval. In DEV (orange) trials, subjects learn that responding earlier in the interval results in a higher overall payoff (higher probability and marginally lower magnitude). **C)** Average behavior on the clock task in Experiment 1 (left panel), the out-of-session Replication session of Experiment 1 (middle panel) and Experiment 2 (right panel). The average response time of subjects in IEV (green) increases, indicating that subjects learn to respond later. During DEV trials (orange) subjects learn to respond earlier in the interval with shorter RTs. Responses in the Average responses in the CEV and CEVR conditions lie in between the IEV and DEV conditions. In Experiment 2, the CEV and CEVR conditions were omitted and there were unsigned reversals between IEV and DEV every 40 trials. **D)** On this task, shifts toward the best-known option (RTVmax) are interpreted as exploitative (right panel) whereas other shifts in RT (left panel) are interpreted as exploration. Note that the left panel has a more complex value distribution (“landscape”) which would result in a higher entropy, whereas the right panel has a single prominent value peak which would result in a lower entropy.

Participants completed eight 50-trial runs of the clock task during an fMRI scan. The clock task was implemented in PsychToolbox 3.0.10 (Brainard, D.H., 1997; Kleiner, M et al., 2007) and run in MATLAB 2012a (The Mathworks, Inc. Natick, Massachusetts, www.mathworks.com). The inter-trial interval varied in length according to an exponential distribution, with the sequence and length optimized for detection power using a Monte Carlo approach implemented by the *optseq2* command in Freesurfer 5.3. Specifically, 5 million possible inter-trial interval sequences were simulated, with the top 320 sequences retained based on their estimation efficiency. For each subject, 8 of these efficient inter-trial interval sequences were selected, one for each scanner run.

Participants also completed 8 runs of the clock task during a separate MEG session (Experiment 1 - non-fMRI Replication Session, or Experiment 1 - Out-of-Session MEG Replication), which was counterbalanced in order with the fMRI session and separated by an average of 3.7 weeks. For the purposes of this paper, this second session allowed examination of stable tendencies to explore or exploit and replication of effects across time. Due to lower signal-to-noise ratio of MEG relative to fMRI, runs consisted of 63 trials.

During each trial, the central stimulus was an abstract phase-scrambled image, or a happy or fearful face. The phase-scrambled images were intended to provide equal luminance and coloration. Faces were selected from the NimStim database (Tottenham et al., 2009). All 4 contingencies were collected with scrambled images, while only IEV and DEV had emotion manipulations. We do not report on central stimulus effects here, presenting average estimates across stimuli.

### BEHAVIORAL TASK – EXPERIMENT 2

Participants completed the clock task described above, but with six modifications. 1) To dissociate value entropy from novelty, the contingency reversed every 40 trials unknown to the participants. 2) Participants completed 240 trials of the clock task in two runs. 3) Only IEV and DEV contingencies were employed. 4) Trials were extended to 5s to accommodate slower psychomotor speed in this older sample. 5) No MEG data were collected in a separate session. 6) The central stimulus was always a generic clock face. Subjects consistently chose longer response times in the IEV condition (green), and shorter response times in the DEV condition (orange) in Experiment 2 (Fig. 2C).

### COMPUTATIONAL MODELING OF BEHAVIOR

To capture learning and value representation on the clock task, we used a previously validated reinforcement learning (RL) model (StrategiC Exploration/Exploitation of Temporal Instrumental Contingencies, or SCEPTIC; Hallquist and Dombrovski, 2019). The SCEPTIC model represents the one-dimensional value landscape of the clock task using a set of unnormalized Gaussian radial basis functions spaced evenly over the interval. Each function has a temporal receptive field with a mean and variance defining its point of maximal sensitivity and the range of times to which it is sensitive. The quantity of interest tracked by the bases is the expected value of a given choice indicated by the response time. To represent time-varying value, the heights of the basis functions are scaled according to a set of *B* weights, *w* = [*w*_1_, *w*_2_, …, *w*_b_]. The contribution of each basis function to the integrated value representation depends on its temporal receptive field:

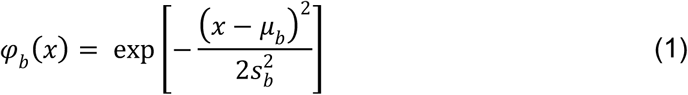

where *x* is an arbitrary point within the time interval *T*, μ_b_ is the center (mean) of the RBF and *s*_b_^2^ is its variance. And more generally, the temporally varying expected value function on a trial *i* is obtained by the multiplication of the weights with the basis:

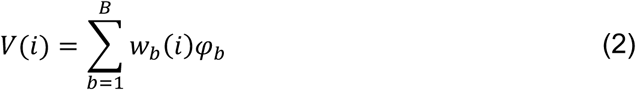

For the clock task, where the probability and magnitude of rewards varied over the course of four-second trials, we spaced the centers of 24 Gaussian RBFs evenly across the discrete interval and chose a fixed width *s*_b_^2^ for the temporal variance (width) of each basis function. More specifically, *s*_b_^2^ was chosen such that the distribution of adjacent radial basis functions overlapped by approximately 50%.

The basic model learns the expected values of different response times by updating each basis function *b* according to the equation:

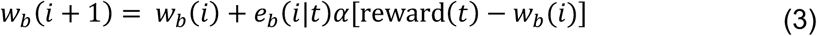

where *i* is the current trial in the task, *t* is the observed response time, and reward(*t*) is the reinforcement obtained on trial *i* given the choice *t*. Prediction error updates are weighted by the learning rate α and the eligibility *e_b_*. To avoid tracking separate value estimates for each possible moment, feedback obtained at a given response time *t* is propagated to adjacent times. This is achieved using a Gaussian RBF centered on the response time *t,* having width *s*_b_^2^. The eligibility *e*_b_ of a basis function φ_b_ to be updated by prediction error is defined as its overlap with the temporal generalization function, *g*:

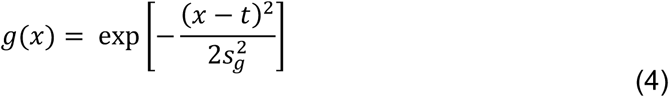

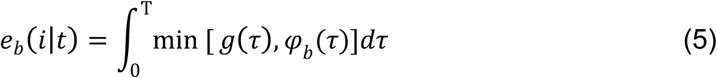

where τ represents an arbitrary timepoint along the interval *T.* Thus, for each radial basis function e*_b_,* a scalar eligibility *e*_b_ between zero and one represents the overlap with the temporal generalization function *s*_b_^2^.

The SCEPTIC model selects an action based on a softmax choice rule, analogous to simpler reinforcement learning problems (e.g., two-armed bandit tasks; Sutton and Barto, 2018). For computational speed, we arbitrarily discretized the interval into 100 ms time bins such that the agent selected among 40 potential responses (i.e., a multinomial representation). At trial *i* the agent chooses a response time in proportion to its expected value:

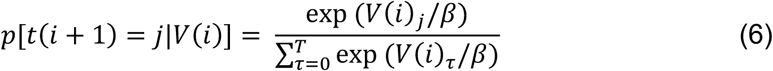

where *j* is a specific response time and the temperature parameter β controls value sensitivity such that choices become more stochastic and less value-sensitive at higher values.

Importantly, as detailed previously (Hallquist and Dombrovski, 2019), selective maintenance of frequently chosen and preferred actions performs better than other alternative models. Specifically, basis weights corresponding to a non-preferred, spatiotemporally distant action revert toward a prior in inverse proportion to the temporal generalization function:

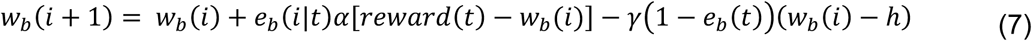

where γ is a selective maintenance parameter between zero and one that scales the amount of decay towards a point *h*, which is taken to be zero here for parsimony, but could be replaced with an alternative prior expectation. Our primary fMRI analyses used signals derived from fitting the information-compressing RL model (eq.7) to participants’ behavior.

We defined the information content of the learned value distribution as Shannon’s entropy of the normalized basis weights (the trial index *i* is omitted for simplicity of notation):

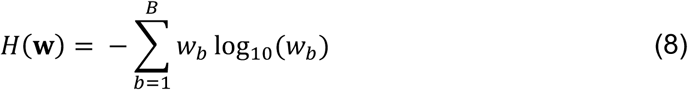

The SCEPTIC model was fit to individual choices using the empirical Variational Bayes Approach (Daunizeau et al., 2014). This method relied on a mixed-effects model where individual-level parameters were assumed to be sampled from a normal distribution.

The group’s summary statistics were then inferred from individual-level posterior parameter estimates using an iterative variational Bayes algorithm that alternates between estimating population parameters and the individual subject parameters. Over algorithm iterations, individual-level priors are minimized toward the inferred parent population distribution, as in standard multilevel regression. The SCEPTIC model was fit separately for Experiment 1 and Experiment 2.

For analysis of fMRI signals trial-by-trial, we used two variables from the SCEPTIC model, the maximum of *V*(*i*), or *V_max_* (Eq. 2), and the entropy *H*(*w*) (Eq. 8).

### FMRI ACQUISITION, PREPROCESSING AND GLM ANALYSIS – EXPERIMENT 1

We used a Siemens Tim Trio 3T scanner with the following T2*-optimized sequence: TR=1.0s, TE=30ms, flip angle = 55°, multiband acceleration factor = 5, voxel size = 2.3mm^3^. T1 anatomical scans for coregistration were acquired with voxel size = 1.0mm^3^, TR=2.2s, TE=3.58ms, and GRAPPA 2x acceleration. A gradient echo fieldmap TE=4.93ms and 7.39ms was acquired to correct for magnetic field inhomogeneity.

Anatomical scans were preprocessed as described previously (Dombrovski et al., 2020). Briefly, slice timing and motion coregistration were performed using NiPy in AFNI (v.19.0.26; Roche, 2011) and FMRIB in FSL (v6.0.1). Skull stripping was performed by masking low intensity voxels and via the ROBEX brain extraction algorithm (Iglesias et al., 2011). Functional images were aligned to T1 anatomical images using white matter segmentation and boundary-based registration (Greve and Fischl, 2009). Given that fast-TR sequences have reduced gray-white contrast, we improved the functional-structural coregistration by first aligning functional volumes to a single-band reference image with improved contrast. Functional scans were then transformed into MNI152 template space (FSL FNIRT) and smoothed using a 5mm FWHM kernel in FSL SUSAN either within the hippocampal mask (for hippocampus signals) or across the whole brain. Head motion artifacts were reduced via an ICA analysis in FSL MELODIC, with the spatiotemporal components then passed to an ICA-AROMA classification algorithm (Pruim et al., 2015). Components that were classified as noise were regressed out of the signal using FSL regfilt (nonaggressive regression approach). Susceptibility artifacts were reduced using fieldmap correction using FSL FUGUE. An 0.008 Hz temporal high-pass filter removed slow-frequency signals (Woolrich et al., 2001). Voxel time series were renormalized to have a mean of 100 to allow similar scaling of voxelwise regression coefficients across run and subject.

Runs where framewise displacement exceeded 10% of total volumes had a framewise displacement of ≥0.9mm, or subjects where peak displacement exceeded 5 mm at any point during the scan were excluded from all analyses. This led to a total of 549 usable runs, with 11 excluded from GLM analysis. There were no subjects for which both runs were excluded due to motion. No runs were excluded for deconvolved signal analyses (described below).

Voxelwise general linear models (GLM) for BOLD data were estimated in FSL (v6.0.1; S. M. Smith et al., 2004) using FSL FEAT (v6.0). All models included regressors for clock and feedback events, calculated by convolving boxcar regressors for clock and feedback events with a double-gamma canonical hemodynamic response function (HRF). In addition, parametric BOLD modulation by either *V_max_* or entropy of the value distribution was modeled by convolving corresponding trial onset-aligned SCEPTIC-derived signals with the HRF. Confound regressors from ventricles and cerebral white matter, as well as derivatives of these signals, were also included. At the second (subject) level, parameter estimates were combined from each run using a weighted fixed effects model in FEAT. Level 3 group analyses were then performed using the FSL FLAME 1+2 approach with automatic outlier deweighting (Woolrich, 2008). All group analyses included age and sex as covariates.

Familywise error correction for voxelwise statistical parametric maps was carried out using Probabilistic Threshold-free Cluster Enhancement method at *p* < 0.05 (Spisák et al., 2019).

### FMRI ACQUISITION AND PREPROCESSING – EXPERIMENT 2

The imaging sessions were conducted at the University of Pittsburgh’s Magnetic Resonance Research Center (RRID:SCR_025215). We used a Siemens Magnetom Prisma scanner with the following T2*-optimized sequence: TR=600ms, TE=27ms, flip angle = 45°, multiband acceleration factor = 5, voxel size = 3.1mm^3^. T1 anatomical scans for coregistration were acquired with voxel size = 1.0mm^3^, TR=2.3s, TE=3.35ms, and GRAPPA 2x acceleration. A gradient echo fieldmap TE=4.47ms and 6.93ms was acquired to correct for magnetic field inhomogeneity.

The fMRI data were then preprocessed using FMRIPREP version 20.1.1 (Roche, 2011), a Nipype-based (Gorgolewski et al., 2011) tool. Each T1w (T1-weighted) volume was corrected for INU (intensity non-uniformity) using N4BiasFieldCorrection v2.1.0 (Tustison et al., 2010) and skull-stripped using antsBrainExtraction.sh v2.1.0 (using the OASIS template). Brain surfaces were reconstructed using recon-all from FreeSurfer v6.0.1 (Dale et al., 1999) and the brain mask estimated previously was refined with a custom variation of the method to reconcile ANTs-derived and FreeSurfer-derived segmentations of the cortical gray-matter of Mindboggle (Klein et al., 2017). Spatial normalization to the ICBM 152 Nonlinear Asymmetrical template version 2009c (Fonov et al., 2009) was performed through nonlinear registration with the antsRegistration tool of ANTs v2.1.0 (Avants et al., 2008) using brain-extracted versions of both T1w volume and template. Brain tissue segmentation of cerebrospinal fluid (CSF), white-matter (WM) and gray-matter (GM) was performed on the brain-extracted T1w using fast FSL v5.0.9 (Zhang et al., 2001).

Functional data was slice-time corrected using 3dTshift from AFNI v16.2.07 (Cox, 1996) and motion corrected using mcflirt (FSL v5.0.9; Jenkinson et al., 2002). Distortion correction was performed using fieldmaps processed with fugue (Jenkinson, 2003).

This was followed by co-registration to the corresponding T1w using boundary-based registration (Greve and Fischl, 2009) with six degrees of freedom, using bbregister (FreeSurfer v6.0.1). Motion correcting transformations, field distortion correcting warp, BOLD-to-T1w transformation and T1w-to-template (MNI) warp were concatenated and applied in a single step using antsApplyTransforms (ANTs v2.1.0) using Lanczos interpolation.

Frame-wise displacement (Power et al., 2014) was calculated for each functional run using the implementation of Nipype. ICA-based Automatic Removal Of Motion Artifacts (AROMA) was used to generate non-aggressive noise regressors (Pruim et al., 2015). Components identified as noise were regressed out of the data using FSL regfilt (non-aggressive regression approach). ICA-AROMA has performed very well in head-to-head comparisons of alternative strategies for reducing head motion artifacts (Ciric et al., 2017). We then applied a .008 Hz temporal high-pass filter to remove slow frequency signal changes (Woolrich et al., 2001); the same filter was applied to all regressors in MEDuSA analyses (see below). Finally, we renormalized each voxel time series to have a mean of 100 to provide similar scaling of voxelwise regression coefficients across runs and participants. Many internal operations of FMRIPREP use Nilearn (Abraham et al., 2014) principally within the BOLD-processing workflow.

### MULTILEVEL EVENT-RELATED DECONVOLVED SIGNAL ANALYSIS (MEDuSA)

The standard GLM approach using canonical HRFs is useful for testing hypotheses about session-level linear associations between model-based signals and regional activity. However, GLMs assume 1) that the correct amplitude and duration of task-related neural signals are specified in the model; and 2) that a canonical HRF describes BOLD activity associated with a model-based signal. In addition, GLMs do not allow the experimenter to probe the time courses of signals before or after trial-level events of interest. Therefore, we employed Multilevel Event-related Deconvolved Signal Analysis (MEDuSA), which operates through voxelwise deconvolution of BOLD activity to functionally align the signal to the event of interest (Dombrovski et al., 2020). In addition, MEDuSA operates within a multilevel framework, providing the experimenter with an appropriate statistical analysis of hierarchically structured data. We briefly outline the MEDuSA pipeline below.

After first applying the preprocessing pipelines described above, a leading hemodynamic deconvolution algorithm was applied voxelwise across the entire run to estimate neural population activity (Bush and Cisler, 2013). Then, the deconvolved signal was extracted from −4s to +4s around trial onset. Linear interpolation was performed to align the deconvolved data onto a TR-consistent grid (1.0s and 0.6s, respectively, for studies 1 and 2). This operation merely resampled the data and did not introduce additional information into the signal. Activity estimates that fell into subsequent trials were censored to ensure analyses only included the time within the current trial. As our behavioral data had a nested structure (trials nested within run, nested within subject), we used multilevel regression models to compute effects.

Multilevel models were estimated using restricted maximum likelihood in the *lme4* package (Bates et al., 2015) in R 4.2.1 (R Core Team, 2021). Estimated *p* values for predictors in the models were computed using Wald chi-square tests and degrees of freedom were based on the Kenward-Roger approximation.

To examine entropy and *V_max_* modulation of vPFC and hippocampus, deconvolved signal in each structure was regressed on entropy and value using a multilevel regression framework. In addition, these analyses included several control variables: prior ITI, current response time, scaled inverted (negative) trial number, outcome, age and gender. Random intercepts of run nested within subject were used, and separate models were computed at each time point in the within-trial grid (−4s to +4s) and each structure. MEDuSA can also be used to estimate functional connectivity between different regions by including one region as an independent variable in a regression analysis predicting activity in another region. Thus, for functional connectivity MEDuSA analyses, multilevel models included vPFC deconvolved signal as the dependent variable and main effects and interactions of within-run change in aligned hippocampal activity and entropy and *V_max_* (as well as several confound regressors) as independent variables. In all analyses, false-discovery rate multiple comparison was applied (Benjamini and Yekutieli, 2001). Additionally, no runs were removed due to framewise displacement, as in the last step of preprocessing for Experiment 1 GLMs.

For the analysis in Fig. 5 and Fig. 6 (Hippocampal signal predicting vPFC activity and Effects on explore/exploit), for both Experiment 1 and 2, activity in the hippocampus was z-scored within subject and run (8 runs in Experiment 1, and 2 runs in Experiment 2) to emphasize relative activity change within a run. The average hippocampal activity for each subject and run was regressed out as a nuisance effect in the multi-level analysis, which controls for BOLD quality effects and conflation of high global signal with the desired functional connectivity signal. The goal of this approach was to capture trial-level fluctuations in hippocampal activity.

To examine how neural activity predicted behavior, the relationship between entropy or and network activity for each participant was extracted from random (participant-varying) effects estimates of the vPFC neural activity models. *x*_H(*w*)_*x*_HC_ case where *HC* is hippocampal activity and *H*(*w*) is entropy from Eq. 8. The outcome term *Y_vPFC_* is computed for each subject *s* and trial *t*. The random slope of the interaction of hippocampal activity with entropy term for each subject is *u*_s_ which gives the subject-level deviation of the regression coefficient from the group regression coefficient γ_0_ (Eq. 9). *u*_s_ *is the key variable we use as a moderator for model-derived effects on behavior*. The overall intercept is γ_0_, and the level 2 regression intercepts are *u*_0s_. The level 1 residual variance is *r*_*st*_.

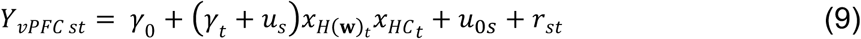

This subject-level random effect from Eq. 9, *u*_s_, was then averaged across the time window and used as a fixed effect to predict upcoming participant response times in a second multilevel model with the lagged response time (exploration term; Eq. 10) or the lagged response time of the *V_max_* (RT*V_max_*, exploitation term). See Fig. 2D for intuition on how correlations between lagged RT with current RT and lagged RT*V_max_* with current RT translate into exploration and exploitation. For clarity that it is a fixed effect predicting *C_RTt_* in Eq. 10, we rename *u*_s_ = *x*_s_. *x*_RT(t-1)t_ is the lagged RT and *x*_R_ is the last outcome, which is a binary variable (reward or omission). The coefficients reported in Fig. 6 represent *C_RTt_* from Eq. 10, estimated *x*_s_ = ±1 standard deviations from the mean using a simple slopes analysis (carried out using the *emtrends* package in R). The significant *p_FDR_* values reported in the text are derived from main effects of the multilevel model.

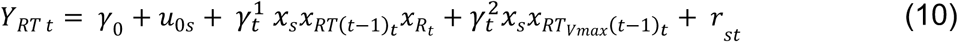

## Results

### VPFC RESPONSE TO ENTROPY OF THE VALUE FUNCTION

Entropy is an emergent property that captures the complexity of the spatially structured value function on a given trial: higher entropy indicates many smaller local value maxima (complex value landscape) and lower entropy typically indicates one prominent global value maximum (simple value landscape; Eq. 8). The higher the value entropy, the more global uncertainty there is about the location of the best option. In exploratory whole-brain voxelwise GLM analyses of Experiment 1 we mapped responses to the entropy of the value function. A cluster responsive to low value entropy in right vPFC (Fig. 3A, black arrow; *p*FDR < 0.05) was centered on the DMN region 24/32, extending into fp10.

**Fig. 3:**
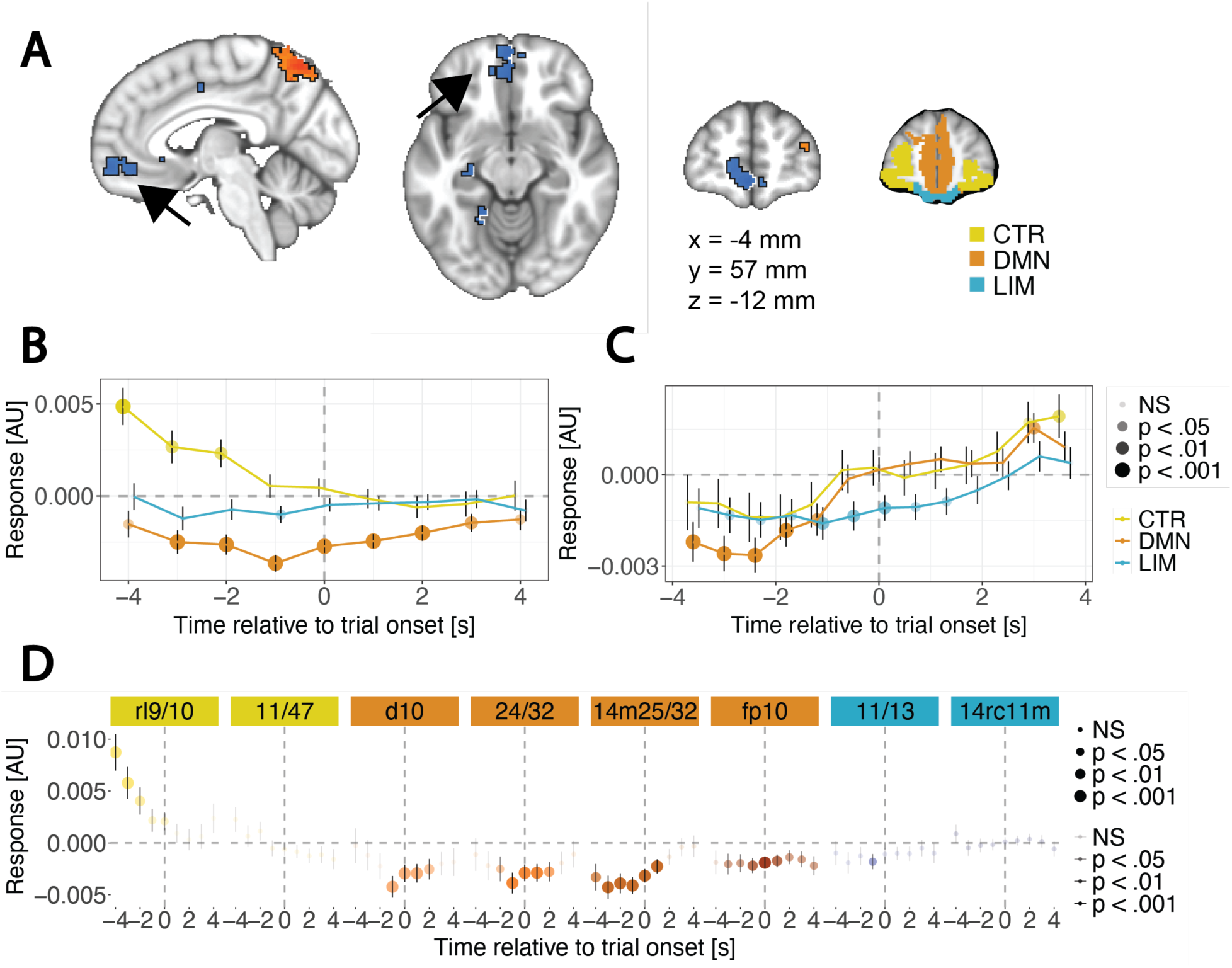
Responses of the vPFC to value function entropy signals. **A.** A whole-brain GLM with a parametric entropy regressor centered on trial onset revealed activation in vPFC (black arrow; DMN region 24/32 extending into fp10). **B.** Entropy was used as a predictor of deconvolved vPFC activity grouped by resting-state network using the MEDuSA method. The DMN (orange) increased activity on lower-value entropy trials, while the CTR (yellow) network increased its activity (FDR-adjusted p-values indicated via opacity and marker size). Timepoints < 0 s represent the inter-trial interval beginning at the offset of previous trial (variable), excluding any volumes from the previous trial. **C.** DMN decreases in response to high entropy (orange) was replicated in Experiment 2. The increased activity of CTR to entropy was observed at a different segment of the window, and an additional strong response to low entropy in LIM was observed. **D.** All regions in the DMN showed response to low value entropy. Only the rl9/10 sub-region of the CTR network increased its activity to higher value entropy.

To investigate regional responses offline (during the inter-trial interval) vs. online (during navigation and choice) we aligned deconvolved BOLD responses to trial onset and examined the within-trial temporal dynamics (Dombrovski et al., 2020). Extending the GLM results, this analysis revealed divergent representations in DMN vs. CTR and no detectable response in LIM (Fig. 3B). DMN subregions responded positively to lower value entropy (orange, Fig. 3B, *p*FDR < 0.001) while CTR subregions responded positively to higher value entropy (yellow, Fig. 3B; *p*FDR < 0.05). In Experiment 2, we replicated the findings of low value entropy responses in DMN (orange, Fig. 3C, 5 points *p*FDR < 0.001) and additionally detected similar responses in LIM (blue, Fig. 3C, *p*FDR < 0.01). High entropy responses in CTR did not replicate in Experiment 2.

We then examined the specific contributions of each of the 17 subregions across the three networks (Fig. 1C) derived from the 200 subregion parcellation of Yeo and Schaefer (Schaefer et al., 2018; see Methods). This analysis of Experiment 1 revealed similar responses to low value entropy across DMN subregions 14m25/32 (orange, Fig. 3D; *p*FDR < 0.05), fp10 (*p*FDR < 0.05), 24/32 (*p*FDR < 0.05) and d10 (*p*FDR < 0.05 between - 1 and 2s). Responses in CTR were more heterogeneous, with dorsal frontopolar region rl9/10 but not orbitofrontal 11/47 responding to higher entropy (yellow, Fig. 3D; 3 timepoints at *p*FDR < 0.05). In LIM, a response to low value entropy was observed only at a single timepoint in region 11/13.

### VPFC RESPONSE TO HIGHEST EXPECTED VALUE (*V_max_*)

While the preceding analysis focused on the global landscape of the value function, scalar expected value signals are commonly reported in vPFC (Hare et al., 2008; Bartra et al., 2013; Smith et al., 2014; Lopez-Gamundi et al., 2021; Knudsen and Wallis, 2022). Thus, to relate our observations to more traditional scalar value analyses of discrete choice tasks, we examined responses to the model-derived expected value of the best option (*V_max_*, Eq. 2, scaled within run to minimize global signal and other run- and subject-level confounds). Our whole-brain voxelwise GLM analysis of Experiment 1 revealed *V_max_* responses in prefrontal DMN (axial slice, region 24/32; Fig. 4A). Significant clusters also included left ventromedial PFC and hippocampus (Table 1).

**Table 1:**
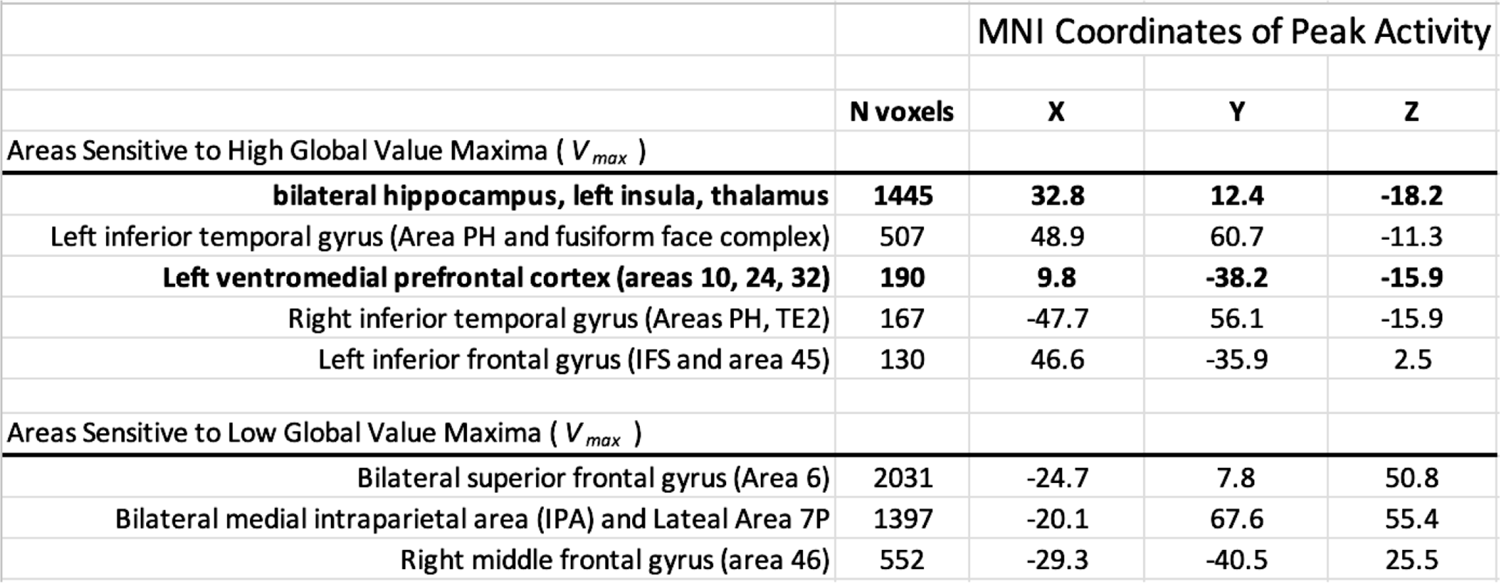
Whole-brain GLM clusters significantly activated by the global value maximum in Experiment 1. Clusters of voxels (voxel size = 2.3mm^3^) significantly modulated by model-derived value maxima (*V_max_*) in whole-brain analyses of Experiment 1. Note: Clusters of significant voxels were identified based on a voxelwise threshold of p < 0.001, with familywise error correction of p < 0.05. Clusters relevant to the present study are in bold.

As above, we followed up by examining within-trial time courses across regions using MEDuSA analyses aligned to trial onset. DMN and LIM vPFC networks responded positively to *V_max_* (Fig. 4B; DMN: orange line, *p*FDR< 0.001; LIM: blue line, *p*FDR< 0.05) and this pattern replicated in Experiment 2 (Fig. 4C). Responses in CTR were absent in Experiment 1 (Fig. 4B, yellow line) and positive in Experiment 2 (Fig. 4C, yellow line).

**Fig. 4:**
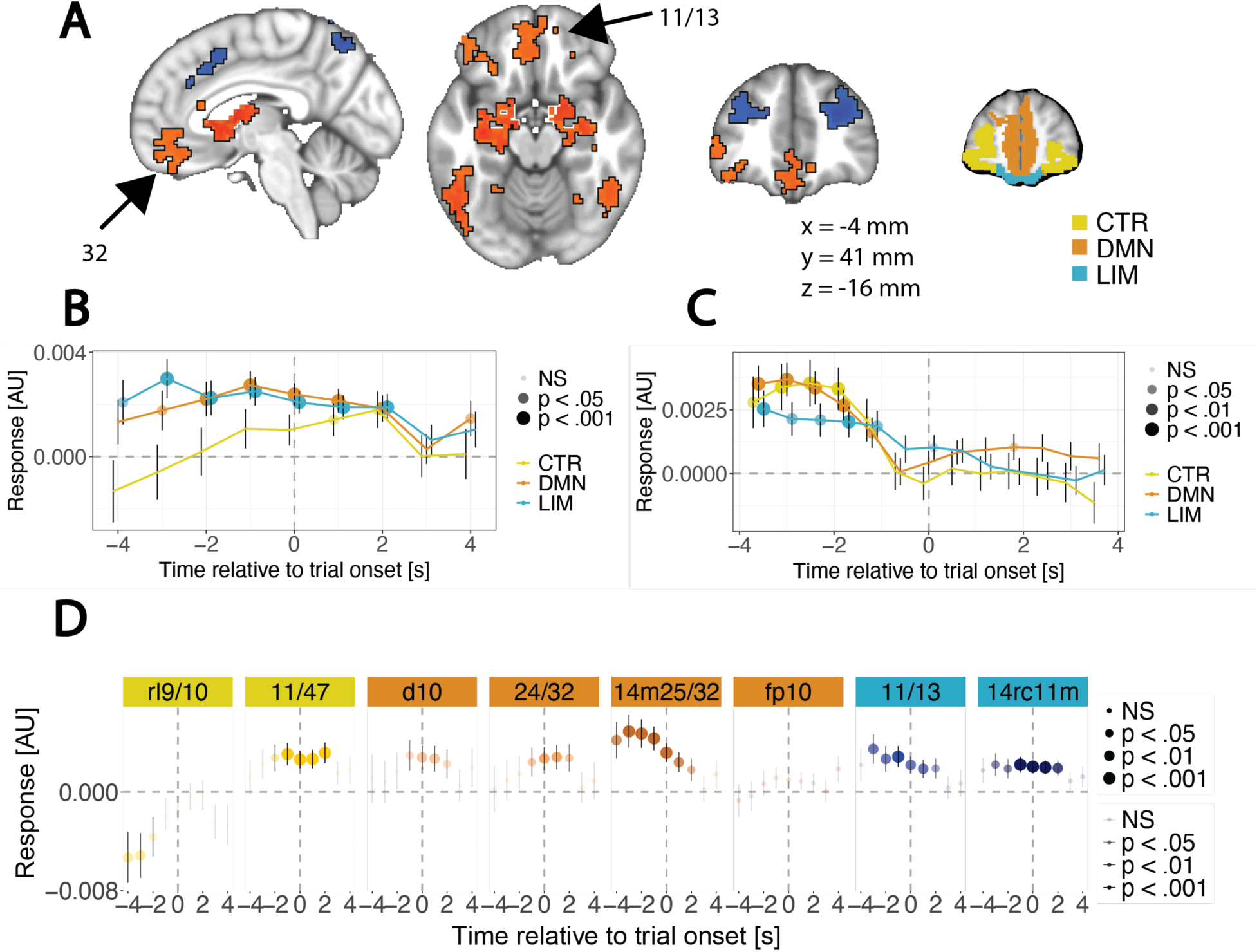
Responses of the vPFC to model-predicted value maximum (*V_max_*) signals. **A.** A whole-brain GLM with a parametric value maximum regressor centered on trial onset revealed significant activation at p_FDR_ < 0.05 in both LIM (axial slice arrow, region 11/13) and DMN network (sagittal slice arrow, multiple regions centered on region 32) visible in the axial slice. **B.** Model-derived *V_max_* was used as a predictor of deconvolved vPFC activity grouped by network using the MEDuSA method. DMN and LIM showed strong responses to scalar value maximum signals. **C.** We replicated our main result in a separate Experiment. DMN (*p*FDR < 0.01) and LIM (*p*FDR < 0.05) increased activity to higher value maxima, but these values occurred earlier in the window than in Experiment 1. **D.** *V_max_* predicted vPFC activity grouped by region in Experiment 1. In the DMN, region 14m25/32 responded strongly with regions 24/32 and d10 responding less prominently. Both LIM regions responded strongly to value signals. In the CTR network, only region 11/47 increased activity to higher *V_max_*, while region rl9/10 increased in response to lower model-derived *V_max_*.

Among vPFC subregions in Experiment 1 (Fig. 5D), DMN 14m25/32 responded most robustly *to V*max (*p*FDR < 0.05). Both LIM subregions 11/3 (*p*FDR < 0.05) and 14rc11m (*p*FDR < 0.05) displayed positive responses. CTR regions displayed divergent responses to *V_max_* with region 11/47 responding to positive signals (*p*FDR < 0.05) and region rl9/10 responding to negative value signals (*p*FDR < 0.05).

**Fig. 5:**
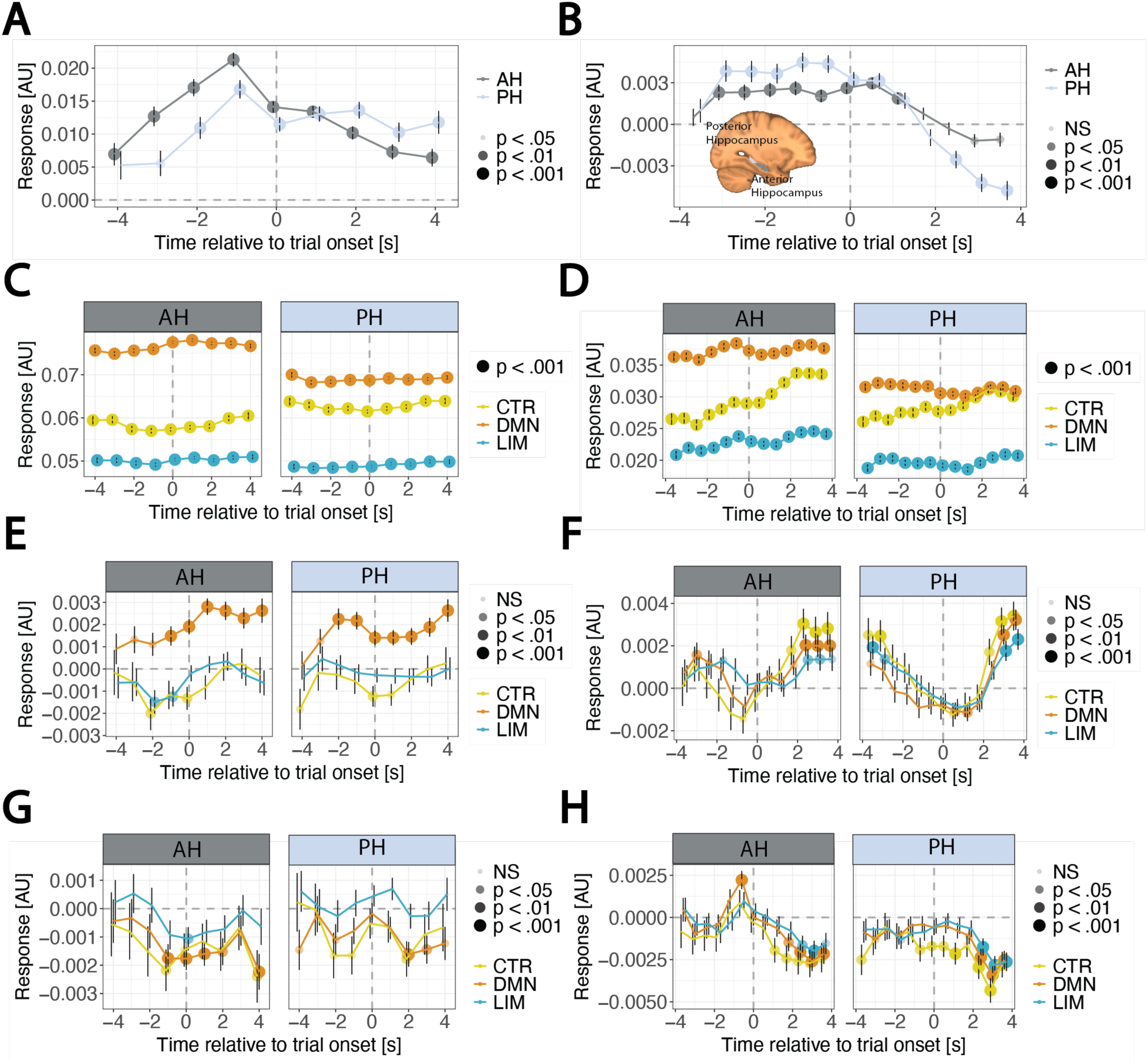
Hippocampus responds to *V_max_* and is selectively synchronized with DMN-vPFC, particularly during periods of higher entropy. **A.** Model-derived *V_max_* was used as a predictor of deconvolved hippocampal activity in anterior (dark grey) and posterior (light gray) hippocampus using the MEDuSA method. Both hippocampal sub-regions responded to *V_max_*. B. *V_max_* responses in hippocampus replicated out-of-sample in Experiment 2. **C.** Hippocampal activity was averaged within region (AH or PH), scaled within subject and run, and then used as a regressor to predict vPFC activity in Experiment 1. AH had higher connectivity with DMN compared to PH. Connectivity generally followed a gradient of DMN > CTR > LIM. **D.** Hippocampal activity was used as a regressor to predict vPFC activity in Experiment 2 following the same procedure. Overall, hippocampal-vPFC connectivity was diminished, but the same gradient of DMN > CTR > LIM remained and AH connectivity was greater than PH connectivity. **E.** Interacting hippocampal signals with entropy for Experiment 1 suggests that DMN is most strongly influenced by hippocampus at times when entropy is higher. **F.** Anterior Hippocampal (AH) activity affects DMN activity more strongly when entropy is higher in Experiment 2 following trial onset, replicating the results out-of-sample. In addition, in this population, hippocampal activity affects CTR more strongly when entropy is higher. **G.** Interacting hippocampal signals with *V_max_* in Experiment 1 suggests that DMN is most strongly influenced by anterior hippocampus at times when *V_max_* is lower. **H.** In Experiment 2, anterior hippocampus influences DMN more strongly during trials when *V_max_* is lower, replicating this finding out-of-sample. In addition, in Experiment 2, the entire vPFC is synched with hippocampus activity at lower *V_max_*, notably during a later time in the window.

### HIPPOCAMPAL VALUE RESPONSE AND HIPPOCAMPAL-vPFC CONNECTIVITY

We previously reported that the hippocampus, particularly its anterior divisions, responded to low-entropy, easy-to-exploit value landscapes, and this response scaled with behavioral exploitation (Dombrovski et al., 2020). Here, we further consider spatiotemporal patterns of hippocampal activity and connectivity with vPFC, including how connectivity is modulated by entropy and *V_max_*, processes that control the explore-exploit balance.

In whole-brain GLM (Table 1) and deconvolved signal analyses aligned to trial onset (Fig. 5A) both the anterior and posterior hippocampus responded to *V_max_* (all p_FDR_ < 0.001). This pattern was strongly replicated in Experiment 2 (Fig. 5B).

To assess functional connectivity, we regressed vPFC activity on contemporaneous hippocampal activity (i.e., at the same moment within the trial). In Experiment 1, AH (dark gray panel) was more strongly functionally coupled to DMN compared to PH (light gray panel), and a clear gradient of hippocampal coupling was observed with DMN > CTR > LIM (Fig. 4C). In Experiment 2, results were qualitatively similar, though functional coupling between hippocampus and vPFC-CTR increased following trial onset (Fig. 6D).

Activity patterns (Fig. 4B-C) led us to expect stronger AH-DMN connectivity during periods of low entropy. Contrary to this expectation, hippocampal-DMN functional coupling was increased during periods of higher entropy in both AH (dark gray) and PH in both experiments (light gray; Fig. 5E-F). Further, in Experiment 2 only, connectivity modulation was seen only during the online portion of the task, post trial onset and both CTR- and LIM-hippocampal connectivity also scaled up with entropy (Fig. 5F). *V_max_* modulated hippocampal connectivity in a similar pattern across both experiments (Fig. 5G-H).

In summary, the highest levels of connectivity with hippocampus were seen in prefrontal DMN regions, in anterior hippocampus, and when decisions were difficult, as indicated by higher entropy or lower *V_max_*.

### HIPPOCAMPAL-vPFC INTERACTIONS AND BEHAVIORAL EXPLORATION

Normatively, high entropy of the value function favors exploration. Surges in hippocampal-prefrontal DMN connectivity during difficult high-entropy and low-value epochs could thus support behavioral exploration, which we previously found to be related to posterior hippocampal learning signals (Dombrovski et al., 2020). Hence, we examined whether value-related increases in PH-prefrontal DMN connectivity were selectively related to exploration. The temporal resolution of fMRI is insufficient for testing momentary connectivity-behavior relationships, so we used session-level estimates of connectivity modulation to predict behavioral exploration both within-session and out-of-session. We derived session-level estimates of connectivity modulation from the models described above (Eq. 9, Fig. 3, Fig. 5) by extracting individual random slopes for the hippocampal response-by-entropy interaction effect on prefrontal activity and averaged them across timepoints within trial to reduce multiple comparisons. These predictors, one for each subject and each region, were then entered into the behavioral model predicting exploration (Eq. 10). As in previous studies, exploration of the continuous space was measured by large departures from a previous response (“RT swings”, Fig. 2D, left panel (Badre et al., 2012; Hallquist and Dombrovski, 2019; Hallquist et al., 2023). While this analysis controlled for long-term value (exploitation) effects, one must also account for the strong win-stay/lose-shift response tendency prominent in humans and other species (Zhang et al., 2023; Randall and Zentall, 1997). Thus, post-reward RT swings provide a purer measure of exploration as defined in RL.

Indeed, stronger modulation of PH-DMN and PH-LIM connectivity by value entropy scaled with exploratory post-reward RT swings (Fig. 6A) in-session (Experiment 1, *p*FDR < 10^-5^), out-of-session (MEG replication, *p*FDR = 0.03) and out-of-sample (Experiment 2, *p*FDR = < 10^-5^). Main effects were significant for LIM-PH in Experiment 1 (*p*FDR = 2.3e-19), Experiment 1 out-of-session MEG replication (*p*FDR = 1.2e-15), and Experiment 2 (*p*FDR = 6.7e-4). Recall that, as Fig. 5E-F illustrates, only PH-DMN, but not PH-LIM connectivity was consistently modulated by entropy across entire samples, qualifying the interpretation of PH-LIM interaction behavioral correlates. Significant post-hoc contrasts are indicated by non-overlapping confidence intervals and consistent effects across all three experimental datasets are indicated by higher opacity. Here, p-values with false discovery rate correction are reported from the entropy connectivity random slope by lagged RT by last outcome term (γ^1^_t_ *x*_s_*x*_RT(t-1)t_ *x_Rt_* in Eq. 10).

To ascertain the anatomical specificity of the relationship between PH-DMN connectivity and exploration, we repeated the above analysis for AH-vPFC connectivity (Fig. 6B).

While modulation of AH-DMN by value entropy was not related to win-shift exploration, it predicted smaller lose-shifts. Further, stronger modulation of AH-LIM connectivity by value entropy predicted win-shift and, to a lesser degree, lose-shift responses.

Modulation of hippocampal-vPFC connectivity by scalar *V_max_* showed largely non-significant exploitation effects (not reported) suggesting altogether that the critical hippocampal-vPFC interactions signal a spatially structured value map and not merely a scalar *V_max_*.

In summary, as suggested by connectivity analyses (Fig. 5E, F), value-related PH-DMN scaled with behavioral exploration. Unexpectedly, the same was true for PH-LIM and AH-LIM interactions.

### HIPPOCAMPAL-vPFC INTERACTIONS AND BEHAVIORAL EXPLOITATION

Our prior findings (Dombrovski et al., 2020) led us to predict that value-related DMN and LIM activity and connectivity with AH would scale with behavioral exploitation. However, we failed to find a replicable general pattern, observing instead only one or the other study correlated with activation. Stronger negative entropy modulation of DMN activity scaled with behavioral exploitation in Experiment 1 (*p*FDR = 0.01, Figure 6C), replicating out-of-session replication (*p*FDR = 0.01), but not out-of-sample (*p*FDR = 0.64). We found no evidence specifically implicating AH vs. PH prefrontal connectivity in exploitation using these analyses. P-values with false discovery rate correction are reported from the entropy connectivity random slope by lagged RT*V_max_* term (γ_t_^2^*x*_s_ *x_RT*V_max_*(t-1)t_* in Eq. 10).

**Fig. 6:**
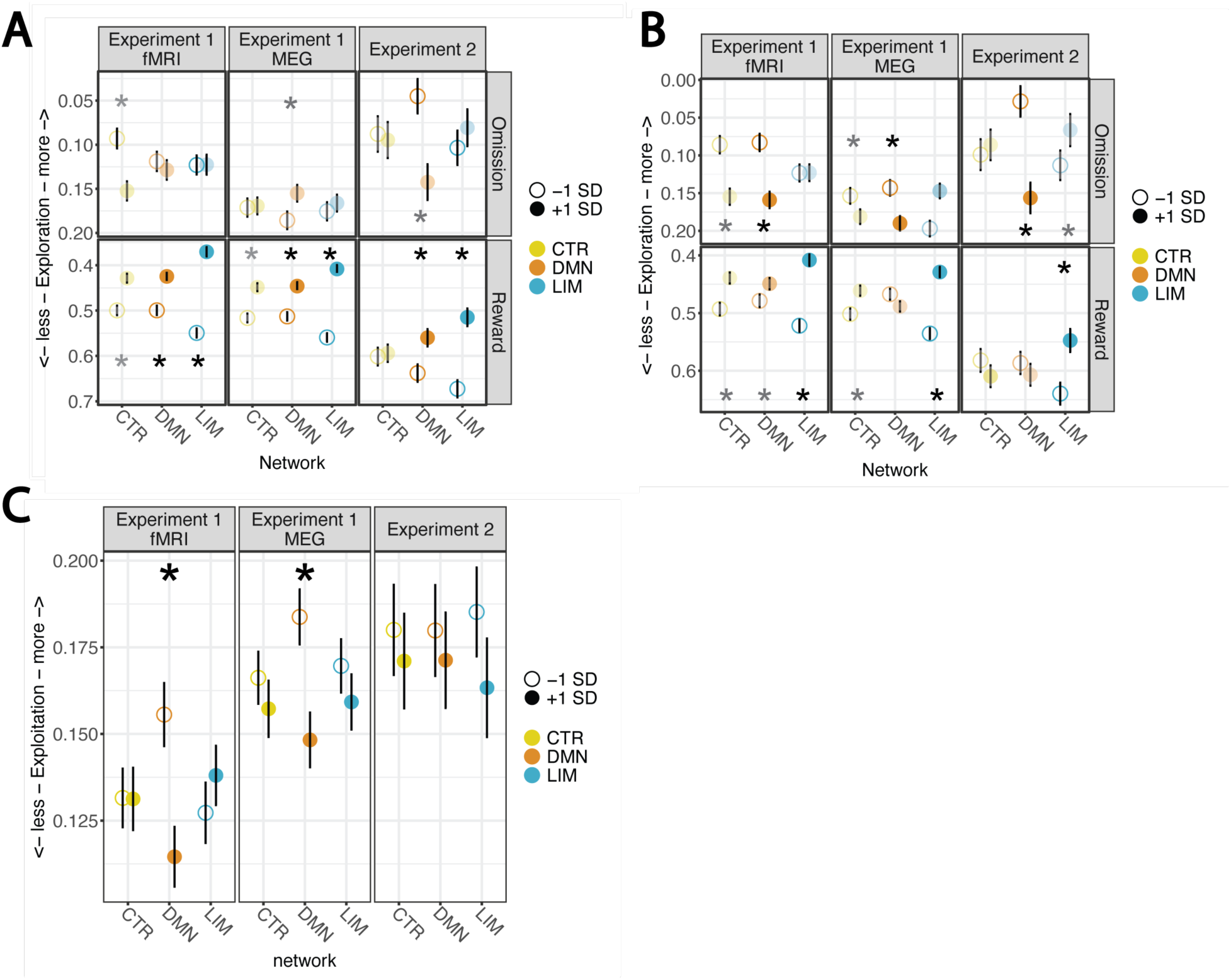
Entropy modulation of vPFC-HC connectivity predicts exploration. The subject-level effects of entropy modulation of vPFC-hippocampus connectivity that predict exploratory behavior of subjects is shown for three datasets. Consistent effect directions across all three datasets are indicated by colored boxes, while significant main effects are indicated by asterisks (*p*FDR reported in text). Higher entropy (one standard deviation above the mean) is indicated by filled colored circles, while lower entropy (one standard deviation below the mean) is indicated by open circles. Note that the y-axes are reversed such that lower autocorrelation between lagged RT and upcoming RT are higher (more Exploration is indicated by lower autocorrelation). For panels A and B, effects that are consistent across all three Experiments are indicated with full opacity, while those that do not replicate are indicated with partial opacity. **A.** When entropy modulation of DMN- and LIM-posterior hippocampal connectivity was higher following reward on the prior trial, subjects engaged in more exploratory behaviors. This win-shift behavior had a significant main effect across all three datasets; thus, we focus on this in the text and discussion. **B.** When entropy modulation of DMN-anterior hippocampus activity was higher following omissions on the prior trial, subjects tended to explore less. This lose-stay behavior was significant across all three datasets. Additionally, when entropy modulation of LIM-anterior hippocampus activity was higher following rewards, subjects engaged in more exploration. This win-shift behavior was significant in Experiment 1 and out-of-session MEG replication session of Experiment 1, but showed a non-significant trend in Experiment 2. **C.** The subject-level effects of entropy modulation of vPFC that predict exploitative behavior of subjects is shown for three datasets. Higher entropy (one standard deviation above the mean) is indicated by filled colored circles, while lower entropy (one standard deviation below the mean) is indicated by open circles. In Experiment 1 and the out-of-session MEG replication session of Experiment 1, lower entropy modulation of DMN predicts exploitation towards the RT*V_max_*.

### SENSITIVITY ANALYSES

Since we assigned the right fp10 region to DMN instead of LIM, we verified that this departure from the previously published parcellation (Thomas Yeo et al., 2011) did not change our results. Reassigning right fp10 back to LIM, findings of DMN activity modulation by low value entropy (Fig. 4B) were qualitatively unchanged (Fig. 7A), as were those for higher *V*max (Fig. 3B, Fig. 7B). Similarly, connectivity patterns reported in Fig. 5C remained qualitatively unchanged (Fig. 7C) and modulation of connectivity by entropy (Fig. 5E) remained qualitatively unchanged (Fig.7D).

**Fig. 7.**
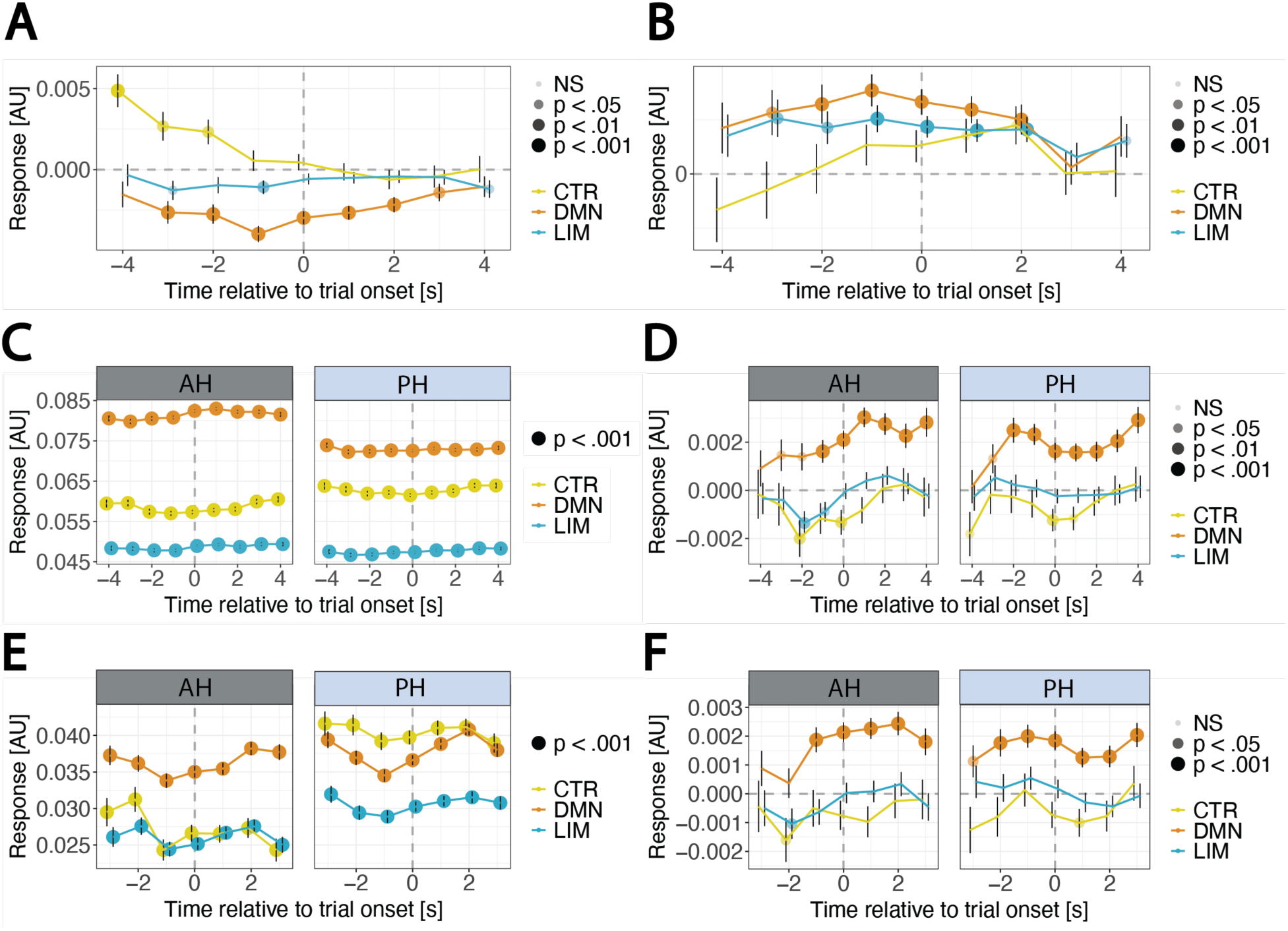
Sensitivity Analyses. In the original 300 parcel, 7 network classification (Schaefer et al., 2018), the region we labeled ‘right fp10’ was assigned to the LIM network instead of the DMN. Here, we check whether re-assigning this parcel to LIM does not alter our major conclusions. **A.** Fig. 3B remains qualitatively unchanged (note a single LIM timepoint at −1s now survives correction at *p*FDR < 0.05) when reassigning right fp10 from DMN to LIM. **B.** Fig. 4B remains qualitatively unchanged (note a slightly smaller effect of *V_max_* on LIM) when reassigning right fp10 from DMN to LIM. **C.** In our functional connectivity model, LIM network correlations with AH were slightly increased and DMN network connectivity with AH was slightly decreased when fp10 was assigned to LIM (compare to Fig. 5C). **D.** Fig. 5E remains qualitatively unchanged after the fp10 parcel is moved to LIM. **E.** Autocorrelation of regressors in time series analysis can lead to spurious significance being detected. To control for this possibility in our vPFC-hippocampal connectivity and psychophysiological interaction analysis, we included the lagged hippocampal signal in our models, controlling for contemporaneous activity of hippocampus. The conclusions of Fig. 5C regarding the gradient of vPFC-hippocampal connectivity are not preserved, but, critically, all points in the connectivity analysis remain significant. **F.** Our finding that hippocampal modulation of DMN is stronger when entropy is higher still holds when accounting for an autocorrelated hippocampal signal, interacting the lagged hippocampal signal with entropy.

Finally, the literature is divided on whether the temporal resolution of fMRI is sufficient for judging the direction of effective connectivity (Barnett and Seth, 2017; Stokes and Purdon, 2017). Still, to ensure that our connectivity results were not explained by temporally slower influences of one region on the other, we repeated our analyses of vPFC-hippocampal connectivity, including both concurrent and *lagged* (by one TR) hippocampal activity. The specific pattern of connectivity reported in Fig. 5C was disrupted (Fig. 7E), but the general significance of connectivity across the time window held. When interacting this divergent pattern of lagged connectivity modulation by value entropy (Fig. 5E vs. Fig. 7F), the general pattern of higher connectivity with DMN during higher value entropy held.

## Discussion

Our study examined the evolving distribution of value across a space modulating vPFC responses and interactions with hippocampus as humans explored and exploited a continuous space. We combined an information-compressing reinforcement learning (RL) model with fMRI to examine how responses to value information (entropy) dynamics in the vPFC and hippocampus shaped the transition from exploration to exploitation. Higher entropy indicates many smaller local value maxima; while lower entropy typically indicates one prominent global value maximum. Entropy representations were more richly captured in pericallosal DMN vPFC subregions than LIM or CTR subregions. Unexpectedly, DMN vPFC-hippocampus functional connectivity increased when people faced uncertain (high entropy, low *V_max_*) value landscapes, and this increase, primarily involving DMN-PH connectivity, scaled with behavioral exploration. These two patterns were observed across sessions, samples and experimental designs. In conclusion, vPFC and hippocampal BOLD activity increases when values are higher and entropy is lower, favoring exploitation. In contrast, hippocampal-vPFC DMN connectivity increases under conditions favoring exploration, when many similarly-valued options compete.

### VALUE REPRESENTATIONS IN NON-HUMAN PRIMATE AND HUMAN VPFC

Why might vPFC subregions of the DMN show greater sensitivity to complex landscapes of learned value? The most straightforward conclusion of our study is that human prefrontal DMN – rather than limbic OFC – contains richer spatial value representations. Monkey lesion and electrophysiological studies of value representations and prefrontal-hippocampal interactions during decision-making mostly converge on the limbic cOFC/mOFC, particularly areas 11, 13 and 14 (Padoa-Schioppa and Assad, 2008; Rolls, 2008; Rudebeck and Murray, 2011; Rich and Wallis, 2014; Pastor-Bernier et al., 2019). By contrast, human imaging studies find value signals in the pericallosal vPFC falling generally into DMN (Kable and Glimcher, 2007; Hare et al., 2008; Bartra et al., 2013; Lim et al., 2013; Vassena et al., 2014; Lopez-Gamundi et al., 2021), which has also become the focus of studies of prefrontal-hippocampal connectivity during decision-making (Gerraty et al., 2014; Gluth et al., 2015; Schuck and Niv, 2019).

Our computational model highlights a related factor potentially responsible for this divergence: whether values are represented at option vs. entire environment level.

While responses to the highest-value option occurred throughout limbic OFC and pericallosal DMN, responses to the entire value landscape were restricted to DMN, particularly area 14m/24/25 and frontopolar dorsal area 10. Both areas, particularly the DMN, synchronized with the hippocampus when value landscapes were complex.

Although human vPFC DMN lacks a direct non-hominoid homologue (Mantini et al., 2011, 2012; Neubert et al., 2015; Garin et al., 2022), our results agree with studies finding preferential medial frontal (Giarrocco and Averbeck, 2021) and DMN (Kaplan et al., 2016) functional connectivity with the hippocampus. Strong scalar value responses in the limbic OFC also dismiss a trivial explanation for the divergence between monkey and human studies—that limbic OFC BOLD is obscured by signal dropout. Indeed, mOFC DMN area 14m/25-32 is equally, if not more, affected by dropout compared e.g. to limbic mOFC area 11/13. Overall, our results suggest that value representations in vPFC DMN, as opposed to limbic OFC, are integrated across the entire environment and more enriched with spatial information from the hippocampus.

### REWARD REPRESENTATIONS WITHIN HIPPOCAMPAL MAPS

Whereas vPFC value representations have been studied in detail, less is known about hippocampal value encoding. Adding to prior evidence of value responses in the human hippocampus (Zeithamova et al., 2018; Bakkour et al., 2019) and our earlier observation that the spatially structured value function selectively modulates AH vs. PH (Dombrovski et al., 2020), here we describe scalar maximum value representations throughout the hippocampal long axis (Fig. 5A-B). In line with our observations, Yun and colleagues (2023) found that rodent dorsal (human posterior) hippocampus responses scaled with discrete-option scalar value and ventral (human anterior) hippocampus represented the maximum value option in a set during a cued licking task. Altogether, hippocampal value representations appear to follow the same fine-coarse postero-anterior (dorso-ventral) gradient as purely spatial representations. If so, one unanswered question is whether this architecture extends to abstract, non-spatial problems. In a recent study, navigation through an abstract space induced value representations in monkey hippocampus (Knudsen and Wallis, 2021), providing evidence of a value-laden cognitive map in a non-spatial context. While the rodent hippocampus is modulated by value-related factors (Wikenheiser and Redish, 2015; Sun et al., 2020), rodent studies have yet to provide evidence that hippocampal maps incorporate value signals in abstract, non-spatial paradigms.

### INTERACTIONS BETWEEN HIPPOCAMPUS AND vPFC

Whereas we anticipated that vPFC and hippocampus would synchronize when people faced easy-to-exploit value landscapes, they instead synchronized when people faced lower values and multiple competing options demanding exploration. Further, while baseline functional connectivity with DMN was stronger in AH than in PH, consistent with resting-state observations (Barnett et al., 2021), it was the synchronization of vPFC-DMN with PH, rather than AH, that scaled with exploratory behavior. This aligns with observations that PH (rodent dorsal hippocampus) represents stimuli during early learning (Biane et al., 2023) and PH/DH learning signals scale with behavioral exploration (Kheirbek et al., 2013; Dombrovski et al., 2020). Further, reward-selective cells in deep DH layers project to the PFC (Harvey et al., 2023), and PH-prefrontal interactions support human memory encoding (Spellman et al., 2015) and retrieval (Rajasethupathy et al., 2015; Ezama et al., 2021).

Although our fMRI analyses do not speak to the electrophysiological mechanisms of PH-prefrontal DMN interactions during exploration, rodent electrophysiological studies offer a basis for speculation. Prefrontal-PH CA1 synchronization involves theta sequences predicting upcoming locations during navigation (“look-ahead”) and sharp wave ripples (SWR) subserving trajectory replay offline, during the inter-trial period (Tang et al., 2021; Wikenheiser and Redish, 2015; Buzsáki, 2015; Norman et al., 2019; Liu et al., 2022). While care should be taken when relating BOLD to electrophysiological responses (Ekstrom et al., 2009; Ekstrom, 2010; Hill et al., 2021), in our analyses of deconvolved BOLD signal value-related PH-DMN interactions extend into the intertrial period. This observation aligns with findings of replay-related interactions between human hippocampus and vPFC interactions (Schuck and Niv, 2019) and broader evidence that key DMN computations and interactions with hippocampus occur offline and during recall (Kaplan et al., 2016; Norman et al., 2021). Thus, when we face multiple competing options, our PH and prefrontal DMN may be jointly simulating exploration trajectories, experienced or contemplated. It is more difficult to interpret the general lack of relationships of value-driven activity and synchronization with exploitation (although in Experiment 1, DMN entropy modulation did predict exploitation; Fig. 6C). It is possible that contributions of hippocampal representations to behavioral exploitation do not depend on coupling with vPFC, but rather on coupling with the dorsal stream network, which we have found to facilitate exploitation on the clock task (Hallquist et al., 2024).

Under the SCEPTIC model, infrequently sampled locations are compressed out of the option set, decreasing value function entropy as one transitions from exploration to exploitation (Hallquist and Dombrovski, 2019). Empirically, vPFC-HC connectivity peaked when entropy was high (Fig. 5E-F), and this modulation scaled with exploratory behavior (Fig. 6A-B). Thus, vPFC-HC coupling may support the compression of an overwhelmingly complex value-laden map to a more manageable and exploitable one. Such plasticity is consistent the broader contributions of the vPFC and hippocampus to behavioral flexibility (Winocur and Olds, 1978; Hornak et al., 2004; Vilà-Balló et al., 2017; McLaughlin and Redish, 2023).

## CONCLUSIONS

Replication across sessions and studies—spanning samples of younger, middle-aged, and older adults, tasks with and without reversals, and scanning sequences— strengthens confidence in our main findings. At the same time, we cannot say which of the differences between studies explain the divergences in the LIM response to entropy (compare Fig. 3B, 3C), timing of *V_max_* signals (compare Fig. 4B, 4C), and timing of entropy modulation of hippocampal-DMN connectivity (compare Fig. 5E, 5F). Without direct manipulation of hippocampal and prefrontal activity, we cannot be confident that hippocampal-vPFC interactions are necessary for behavioral exploration, despite controlling for behavioral confounds, inter-trial interval, age, and sex. The observational study design and low temporal resolution of fMRI make it hard to determine the direction of information transfer during hippocampal-vPFC interactions.

In summary, vPFC and hippocampal BOLD activity increases when the values of dominant options are high, favoring exploitation. Conversely, hippocampal-vPFC DMN connectivity rises under precisely the opposite conditions, when people explore multiple competing options of relatively low value. Coordinated activity of the posterior hippocampal-prefrontal DMN system updates detailed value-laden maps that guide exploration. Given the connection between exploration in the laboratory and human psychopathology, including suicidal behavior (Tsypes et al., 2024), our findings may also inform our understanding of the role of PH-DMN and AH-LIM interactions in real-life decisions at longer timescales and with higher stakes.

## Acknowledgements

This work was funded by K01 MH097091, R01 MH10095, K23 MH122626 and R01 MH067924 from the National Institute of Mental Health. This research was supported in part by the University of Pittsburgh Center for Research Computing through the resources provided. Specifically, this work used the H2P cluster, which is supported by NSF award number OAC-2117681 and the HTC cluster, which is supported by NIH award number S10OD028483. This research was also performed, in part, using resources and the computing assistance of the Pennsylvania State University Institute for CyberScience Advanced CyberInfrastructure (ICS-ACI). We would like to thank Jiazhou Chen, Morgan Buerke, Mandy Collier, Michelle Perry, Laura Taglioni, Shreya Sheth, Tanya Shah, Kaylee Stewart, and Bea Langer for collecting the fMRI data in Experiment 2. We would like to thank Kai Hwang and Rajpreet Chahal for collecting the fMRI data in Experiment 1. We would like to thank Katalin Szanto for administrating the longitudinal study of which Experiment 2 was the scanning arm. Thanks to Nathaniel J. Powell for comments on this manuscript.

## References

Abraham A, Pedregosa F, Eickenberg M, Gervais P, Mueller A, Kossaifi J, Gramfort A, Thirion B, Varoquaux G (2014) Machine learning for neuroimaging with scikit-learn. Front Neuroinformatics 8 Available at: https://www.frontiersin.org/articles/10.3389/fninf.2014.00014 [Accessed May 5, 2023].

Badre D, Doll BB, Long NM, Frank MJ (2012) Rostrolateral prefrontal cortex and individual differences in uncertainty-driven exploration. Neuron 73:595–607.

Bakkour A, Palombo DJ, Zylberberg A, Kang YH, Reid A, Verfaellie M, Shadlen MN, Shohamy D (2019) The hippocampus supports deliberation during value-based decisions Kahnt T, Frank MJ, Fellows LK, Gluth S, eds. eLife 8:e46080.

Barnett AJ, Reilly W, Dimsdale-Zucker HR, Mizrak E, Reagh Z, Ranganath C (2021) Intrinsic connectivity reveals functionally distinct cortico-hippocampal networks in the human brain. PLOS Biol 19:e3001275.

Barnett L, Seth AK (2017) Detectability of Granger causality for subsampled continuous-time neurophysiological processes. J Neurosci Methods 275:93–121.

Bartra O, McGuire JT, Kable JW (2013) The valuation system: a coordinate-based meta-analysis of BOLD fMRI experiments examining neural correlates of subjective value. Neuroimage 76:412–427.

Bates D, Mächler M, Bolker B, Walker S (2015) Fitting Linear Mixed-Effects Models Using lme4. J Stat Softw 67:1–48.

Benjamini Y, Yekutieli D (2001) The control of the false discovery rate in multiple testing under dependency. Ann Stat 29:1165–1188.

Biane JS, Ladow MA, Stefanini F, Boddu SP, Fan A, Hassan S, Dundar N, Apodaca-Montano DL, Zhou LZ, Fayner V, Woods NI, Kheirbek MA (2023) Neural dynamics underlying associative learning in the dorsal and ventral hippocampus. Nat Neurosci:1–12.

Brainard, D.H. (1997) The Psychophysics Toolbox. Spat Vis 10:433–436.

Bush K, Cisler J (2013) Decoding neural events from fMRI BOLD signal: A comparison of existing approaches and development of a new algorithm. Magn Reson Imaging 31:976–989.

Buzsáki, G. (2006) Rhythms in the Brain. Oxford University Press, USA. Available at: https://academic.oup.com/book/11166.

Buzsáki G (2015) Hippocampal sharp wave-ripple: A cognitive biomarker for episodic memory and planning. Hippocampus 25:1073–1188.

Chudasama Y, Doobay VM, Liu Y (2012) Hippocampal-Prefrontal Cortical Circuit Mediates Inhibitory Response Control in the Rat. J Neurosci 32:10915–10924.

Costa VD, Mitz AR, Averbeck BB (2019) Subcortical Substrates of Explore-Exploit Decisions in Primates. Neuron 103:533–545.e5.

Cox RW (1996) AFNI: software for analysis and visualization of functional magnetic resonance neuroimages. Comput Biomed Res 29:162–173.

Dale AM, Fischl B, Sereno MI (1999) Cortical Surface-Based Analysis: I. Segmentation and Surface Reconstruction. NeuroImage 9:179–194.

Daunizeau J, Adam V, Rigoux L (2014) VBA: A Probabilistic Treatment of Nonlinear Models for Neurobiological and Behavioural Data. PLOS Comput Biol 10:e1003441.

Dombrovski AY, Luna B, Hallquist MN (2020) Differential reinforcement encoding along the hippocampal long axis helps resolve the explore–exploit dilemma. Nat Commun 11:5407.

Domenech P, Rheims S, Koechlin E (2020) Neural mechanisms resolving exploitation-exploration dilemmas in the medial prefrontal cortex. Science 369:eabb0184.

Eickhoff SB, Paus T, Caspers S, Grosbras M-H, Evans AC, Zilles K, Amunts K (2007) Assignment of functional activations to probabilistic cytoarchitectonic areas revisited. NeuroImage 36:511–521.

Eickhoff SB, Stephan KE, Mohlberg H, Grefkes C, Fink GR, Amunts K, Zilles K (2005) A new SPM toolbox for combining probabilistic cytoarchitectonic maps and functional imaging data. NeuroImage 25:1325–1335.

Ekstrom A (2010) How and when the fMRI BOLD signal relates to underlying neural activity: The danger in dissociation. Brain Res Rev 62:233–244.

Ekstrom A, Suthana N, Millett D, Fried I, Bookheimer S (2009) Correlation Between BOLD fMRI and Theta-Band Local Field Potentials in the Human Hippocampal Area. J Neurophysiol 101:2668–2678.

Ezama L, Hernández-Cabrera JA, Seoane S, Pereda E, Janssen N (2021) Functional connectivity of the hippocampus and its subfields in resting-state networks. Eur J Neurosci 53:3378–3393.

Fonov V, Evans A, McKinstry R, Almli C, Collins D (2009) Unbiased nonlinear average age-appropriate brain templates from birth to adulthood. NeuroImage 47:S102.

Garin CM, Hori Y, Everling S, Whitlow CT, Calabro FJ, Luna B, Froesel M, Gacoin M, Ben Hamed S, Dhenain M, Constantinidis C (2022) An evolutionary gap in primate default mode network organization. Cell Rep 39:110669.

Gerraty RT, Davidow JY, Wimmer GE, Kahn I, Shohamy D (2014) Transfer of Learning Relates to Intrinsic Connectivity between Hippocampus, Ventromedial Prefrontal Cortex, and Large-Scale Networks. J Neurosci 34:11297–11303.

Giarrocco F, Averbeck BB (2021) Organization of parietoprefrontal and temporoprefrontal networks in the macaque. J Neurophysiol 126:1289–1309.

Gluth S, Sommer T, Rieskamp J, Büchel C (2015) Effective Connectivity between Hippocampus and Ventromedial Prefrontal Cortex Controls Preferential Choices from Memory. Neuron 86:1078–1090.

Gorgolewski K, Burns C, Madison C, Clark D, Halchenko Y, Waskom M, Ghosh S (2011) Nipype: A Flexible, Lightweight and Extensible Neuroimaging Data Processing Framework in Python. Front Neuroinformatics 5 Available at: https://www.frontiersin.org/articles/10.3389/fninf.2011.00013 [Accessed January 23, 2023].

Greve DN, Fischl B (2009) Accurate and robust brain image alignment using boundary-based registration. NeuroImage 48:63–72.

Hallquist M, Hwang K, Luna B, Dombrovski AY (2023) Reinforcement-based option competition in human dorsal stream during exploration/exploitation of a continuous space.: 2023.05.22.541828 Available at: https://www.biorxiv.org/content/10.1101/2023.05.22.541828v1 [Accessed May 23, 2023].

Hallquist MN, Dombrovski AY (2019) Selective maintenance of value information helps resolve the exploration/exploitation dilemma. Cognition 183:226–243.

Hallquist MN, Hwang K, Luna B, Dombrovski AY (2024) Reward-based option competition in human dorsal stream and transition from stochastic exploration to exploitation in continuous space. Sci Adv.

Hare TA, O’Doherty J, Camerer CF, Schultz W, Rangel A (2008) Dissociating the Role of the Orbitofrontal Cortex and the Striatum in the Computation of Goal Values and Prediction Errors. J Neurosci 28:5623–5630.

Harvey RE, Robinson HL, Liu C, Oliva A, Fernandez-Ruiz A (2023) Hippocampo-cortical circuits for selective memory encoding, routing, and replay. Neuron:S0896627323003008.

Henssen A, Zilles K, Palomero-Gallagher N, Schleicher A, Mohlberg H, Gerboga F, Eickhoff SB, Bludau S, Amunts K (2016) Cytoarchitecture and probability maps of the human medial orbitofrontal cortex. Cortex 75:87–112.

Hill PF, Seger SE, Yoo HB, King DR, Wang DX, Lega BC, Rugg MD (2021) Distinct Neurophysiological Correlates of the fMRI BOLD Signal in the Hippocampus and Neocortex. J Neurosci 41:6343–6352.

Hornak J, O’Doherty J, Bramham J, Rolls ET, Morris RG, Bullock PR, Polkey CE (2004) Reward-related Reversal Learning after Surgical Excisions in Orbito-frontal or Dorsolateral Prefrontal Cortex in Humans. J Cogn Neurosci 16:463–478.

Jenkinson M (2003) Fast, automated, N-dimensional phase-unwrapping algorithm. Magn Reson Med 49:193–197.

Jenkinson M, Bannister P, Brady M, Smith S (2002) Improved Optimization for the Robust and Accurate Linear Registration and Motion Correction of Brain Images. NeuroImage 17:825–841.

Johnson A, Redish AD (2007) Neural Ensembles in CA3 Transiently Encode Paths Forward of the Animal at a Decision Point. J Neurosci 27:12176–12189.

Johnson A, Varberg Z, Benhardus J, Maahs A, Schrater P (2012) The hippocampus and exploration: dynamically evolving behavior and neural representations. Front Hum Neurosci 6 Available at: https://www.frontiersin.org/articles/10.3389/fnhum.2012.00216 [Accessed February 7, 2024].

Jones MW, Wilson MA (2005) Theta Rhythms Coordinate Hippocampal–Prefrontal Interactions in a Spatial Memory Task. PLoS Biol 3:e402.

Joo HR, Frank LM (2018) The hippocampal sharp wave-ripple in memory retrieval for immediate use and consolidation. Nat Rev Neurosci 19:744–757.

Kable JW, Glimcher PW (2007) The neural correlates of subjective value during intertemporal choice. Nat Neurosci 10:1625–1633.

Kaplan R, Adhikari MH, Hindriks R, Mantini D, Murayama Y, Logothetis NK, Deco G (2016) Hippocampal Sharp-Wave Ripples Influence Selective Activation of the Default Mode Network. Curr Biol 26:686–691.

Kheirbek MA, Drew LJ, Burghardt NS, Costantini DO, Tannenholz L, Ahmari SE, Zeng H, Fenton AA, Hen R (2013) Differential Control of Learning and Anxiety along the Dorsoventral Axis of the Dentate Gyrus. Neuron 77:955–968.

Klein A, Ghosh SS, Bao FS, Giard J, Häme Y, Stavsky E, Lee N, Rossa B, Reuter M, Neto EC, Keshavan A (2017) Mindboggling morphometry of human brains. PLOS Comput Biol 13:e1005350.

Kleiner, M, Brainard, D.H., Pelli, D., Ingling, A., Murray, R., Broussard, C. (2007) What’s new in Psychtoolbox-3. Perception 36:1–16.

Knudsen EB, Wallis JD (2021) Hippocampal neurons construct a map of an abstract value space. Cell 184:4640–4650.e10.

Knudsen EB, Wallis JD (2022) Taking stock of value in the orbitofrontal cortex. Nat Rev Neurosci 23:428–438.

Laureiro-Martínez D, Brusoni S, Canessa N, Zollo M (2015) Understanding the exploration–exploitation dilemma: An fMRI study of attention control and decision-making performance. Strateg Manag J 36:319–338.

Lim S-L, O’Doherty JP, Rangel A (2013) Stimulus Value Signals in Ventromedial PFC Reflect the Integration of Attribute Value Signals Computed in Fusiform Gyrus and Posterior Superior Temporal Gyrus. J Neurosci 33:8729–8741.

Liu AA et al. (2022) A consensus statement on detection of hippocampal sharp wave ripples and differentiation from other fast oscillations. Nat Commun 13:6000.

Liu Y, Dolan RJ, Kurth-Nelson Z, Behrens TEJ (2019) Human Replay Spontaneously Reorganizes Experience. Cell 178:640–652.e14.

Lopez-Gamundi P, Yao Y-W, Chong TT-J, Heekeren HR, Mas-Herrero E, Marco-Pallarés J (2021) The neural basis of effort valuation: A meta-analysis of functional magnetic resonance imaging studies. Neurosci Biobehav Rev 131:1275–1287.

Lu H, Zou Q, Gu H, Raichle ME, Stein EA, Yang Y (2012) Rat brains also have a default mode network. Proc Natl Acad Sci 109:3979–3984.

Mantini D, Corbetta M, Romani GL, Orban GA, Vanduffel W (2012) Data-driven analysis of analogous brain networks in monkeys and humans during natural vision. NeuroImage 63:1107–1118.

Mantini D, Gerits A, Nelissen K, Durand J-B, Joly O, Simone L, Sawamura H, Wardak C, Orban GA, Buckner RL, Vanduffel W (2011) Default Mode of Brain Function in Monkeys. J Neurosci 31:12954–12962.

McLaughlin AE, Redish AD (2023) Optogenetic disruption of the prelimbic cortex alters long-term decision strategy but not valuation on a spatial delay discounting task. Neurobiol Learn Mem 200:107734.

Mehlhorn K, Newell BR, Todd PM, Lee MD, Morgan K, Braithwaite VA, Hausmann D, Fiedler K, Gonzalez C (2015) Unpacking the exploration–exploitation tradeoff: A synthesis of human and animal literatures. Decision 2:191–215.

Morici JF, Weisstaub NV, Zold CL (2022) Hippocampal-medial prefrontal cortex network dynamics predict performance during retrieval in a context-guided object memory task. Proc Natl Acad Sci 119:e2203024119.

Moustafa AA, Cohen MX, Sherman SJ, Frank MJ (2008) A role for dopamine in temporal decision making and reward maximization in Parkinsonism. J Neurosci 28:12294–12304.

Neubert F-X, Mars RB, Sallet J, Rushworth MFS (2015) Connectivity reveals relationship of brain areas for reward-guided learning and decision making in human and monkey frontal cortex. Proc Natl Acad Sci 112:E2695–E2704.

Norman Y, Raccah O, Liu S, Parvizi J, Malach R (2021) Hippocampal ripples and their coordinated dialogue with the default mode network during recent and remote recollection. Neuron 109:2767–2780.e5.

Norman Y, Yeagle EM, Khuvis S, Harel M, Mehta AD, Malach R (2019) Hippocampal sharp-wave ripples linked to visual episodic recollection in humans. Science 365:eaax1030.

Padoa-Schioppa C (2007) Orbitofrontal cortex and the computation of economic value. Ann N Acad Sci 1121:232–253.

Padoa-Schioppa C, Assad JA (2008) The representation of economic value in the orbitofrontal cortex is invariant for changes of menu. Nat Neurosci 11:95–102.

Papale AE, Stott JJ, Powell NJ, Regier PS, Redish AD (2012) Interactions between deliberation and delay-discounting in rats. Cogn Affect Behav Neurosci 12:513– 526.

Pastor-Bernier A, Stasiak A, Schultz W (2019) Orbitofrontal signals for two-component choice options comply with indifference curves of Revealed Preference Theory. Nat Commun 10:4885.

Power JD, Mitra A, Laumann TO, Snyder AZ, Schlaggar BL, Petersen SE (2014) Methods to detect, characterize, and remove motion artifact in resting state fMRI. NeuroImage 84:320–341.

Pruim RHR, Mennes M, van Rooij D, Llera A, Buitelaar JK, Beckmann CF (2015) ICA-AROMA: A robust ICA-based strategy for removing motion artifacts from fMRI data. NeuroImage 112:267–277.

R Core Team (2021) R: A language and environment for statistical computing. R Foundation for Statistical Computing, Vienna, Austria. Available at: https://www.R-project.org/.

Rajasethupathy P, Sankaran S, Marshel JH, Kim CK, Ferenczi E, Lee SY, Berndt A, Ramakrishnan C, Jaffe A, Lo M, Liston C, Deisseroth K (2015) Projections from neocortex mediate top-down control of memory retrieval. Nature 526:653–659.

Randall CK, Zentall TR (1997) Win-stay/lose-shift and win-shift/lose-stay learning by pigeons in the absence of overt response mediation. Behav Processes 41:227– 236.

Redish AD (2016) Vicarious trial and error. Nat Rev Neurosci 17:147–159.

Rich EL, Wallis JD (2014) Medial-lateral Organization of the Orbitofrontal Cortex. J Cogn Neurosci 26:1347–1362.

Rolls E (2008) Functions of the orbitofrontal and pregenual cingulate cortex in taste, olfaction, appetite and emotion. Acta Physiol Hung 95:131–164.

Royer S, Sirota A, Patel J, Buzsáki G (2010) Distinct Representations and Theta Dynamics in Dorsal and Ventral Hippocampus. J Neurosci 30:1777–1787.

Rudebeck PH, Murray EA (2011) Dissociable Effects of Subtotal Lesions within the Macaque Orbital Prefrontal Cortex on Reward-Guided Behavior. J Neurosci 31:10569–10578.

Schaefer A, Kong R, Gordon EM, Laumann TO, Zuo X-N, Holmes AJ, Eickhoff SB, Yeo BTT (2018) Local-Global Parcellation of the Human Cerebral Cortex from Intrinsic Functional Connectivity MRI. Cereb Cortex 28:3095–3114.

Schmidt B, Redish AD (2021) Disrupting the medial prefrontal cortex with DREADDs alters hippocampal sharp-wave ripples and their associated cognitive processes. Hippocampus 31:1051–1067.

Schuck NW, Niv Y (2019) Sequential replay of nonspatial task states in the human hippocampus. Science 364:eaaw5181.

Schultheiss NW, Schlecht M, Jayachandran M, Brooks DR, McGlothan JL, Guilarte TR, Allen TA (2020) Awake delta and theta-rhythmic hippocampal network modes during intermittent locomotor behaviors in the rat. Behav Neurosci 134:529–546.

Skaggs WE, McNaughton BL (1996) Replay of Neuronal Firing Sequences in Rat Hippocampus During Sleep Following Spatial Experience. Science 271:1870– 1873.

Smith DV, Clithero JA, Boltuck SE, Huettel SA (2014) Functional connectivity with ventromedial prefrontal cortex reflects subjective value for social rewards. Soc Cogn Affect Neurosci 9:2017–2025.

Spellman T, Rigotti M, Ahmari SE, Fusi S, Gogos JA, Gordon JA (2015) Hippocampal– prefrontal input supports spatial encoding in working memory. Nature 522:309– 314.

Stachenfeld KL, Botvinick MM, Gershman SJ (2017) The hippocampus as a predictive map. Nat Neurosci 20:1643–1653.

Stokes PA, Purdon PL (2017) A study of problems encountered in Granger causality analysis from a neuroscience perspective. Proc Natl Acad Sci 114 Available at: https://pnas.org/doi/full/10.1073/pnas.1704663114 [Accessed March 21, 2023].

Sun C, Yang W, Martin J, Tonegawa S (2020) Hippocampal neurons represent events as transferable units of experience. Nat Neurosci 23:651–663.

Sutton RS, Barto AG (2018) Reinforcement Learning, second edition: An Introduction. MIT Press.

Tang W, Jadhav SP (2019) Sharp-wave ripples as a signature of hippocampal-prefrontal reactivation for memory during sleep and waking states. Neurobiol Learn Mem 160:11–20.

Tang W, Shin JD, Jadhav SP (2021) Multiple time-scales of decision-making in the hippocampus and prefrontal cortex. eLife 10:e66227.

Tottenham N, Tanaka JW, Leon AC, McCarry T, Nurse M, Hare TA, Marcus DJ, Westerlund A, Casey B, Nelson C (2009) The NimStim set of facial expressions: judgments from untrained research participants. Psychiatry Res 168:242–249.

Trudel N, Scholl J, Klein-Flügge MC, Fouragnan E, Tankelevitch L, Wittmann MK, Rushworth MFS (2021) Polarity of uncertainty representation during exploration and exploitation in ventromedial prefrontal cortex. Nat Hum Behav 5:83–98.

Tsypes A, Hallquist MN, Ianni A, Kaurin A, Wright AGC, Dombrovski AY (2024) Exploration-Exploitation and Suicidal Behavior in Borderline Personality Disorder and Depression. JAMA Psychiatry Available at: 10.1001/jamapsychiatry.2024.1796 [Accessed August 6, 2024].

Tustison NJ, Avants BB, Cook PA, Zheng Y, Egan A, Yushkevich PA, Gee JC (2010) N4ITK: Improved N3 Bias Correction. IEEE Trans Med Imaging 29:1310–1320.

Ursu S, Clark KA, Stenger VA, Carter CS (2008) Distinguishing expected negative outcomes from preparatory control in the human orbitofrontal cortex. Brain Res 1227:110–119.

Vassena E, Krebs RM, Silvetti M, Fias W, Verguts T (2014) Dissociating contributions of ACC and vmPFC in reward prediction, outcome, and choice. Neuropsychologia 59:112–123.

Vilà-Balló A, Mas-Herrero E, Ripollés P, Simó M, Miró J, Cucurell D, López-Barroso D, Juncadella M, Marco-Pallarés J, Falip M, Rodríguez-Fornells A (2017) Unraveling the Role of the Hippocampus in Reversal Learning. J Neurosci Off J Soc Neurosci 37:6686–6697.

Weilbächer RA, Gluth S (2016) The Interplay of Hippocampus and Ventromedial Prefrontal Cortex in Memory-Based Decision Making. Brain Sci 7:4.

Wikenheiser AM, Redish AD (2015) Hippocampal theta sequences reflect current goals. Nat Neurosci 18:289–294.

Winocur G, Olds J (1978) Effects of context manipulation on memory and reversal learning in rats with hippocampal lesions. J Comp Physiol Psychol 92:312–321.

Woolrich MW, Ripley BD, Brady M, Smith SM (2001) Temporal Autocorrelation in Univariate Linear Modeling of FMRI Data. NeuroImage 14:1370–1386.

Yeo BTT, Krienen FM, Jorge Sabuncu MR, Lashkari D, Hollinshead M, Roffman JL, Smoller JW, Zöllei L, Polimeni JR, Fischl B, Liu H, Buckner RL (2011) The organization of the human cerebral cortex estimated by intrinsic functional connectivity. J Neurophysiol 106:1125–1165.

Yun M, Hwang JY, Jung MW (2023) Septotemporal variations in hippocampal value and outcome processing. Cell Rep 42:112094.

Zeithamova D, Gelman BD, Frank L, Preston AR (2018) Abstract Representation of Prospective Reward in the Hippocampus. J Neurosci 38:10093–10101.

Zhang Y, Brady M, Smith S (2001) Segmentation of brain MR images through a hidden Markov random field model and the expectation-maximization algorithm. IEEE Trans Med Imaging 20:45–57.

Zhang Y, Huynh TKT, Dyson BJ (2023) Deliberately making miskates: Behavioural consistency under win maximization and loss maximization conditions. Npj Sci Learn 8:1–7.

